# Multidimensional topography of memory revealed from thousands of daily documented memories

**DOI:** 10.1101/2022.07.29.501921

**Authors:** Wilma A. Bainbridge, Chris I. Baker

## Abstract

Our memories form a rich, colorful tapestry of emotions, events, people, and places, woven across the decades of our lives. However, research has typically been limited in its ability to assess the multidimensional nature of episodic memory, given the short time scales and artificial stimulus sets often required in experiments. In an era when people are constantly recording their lives through social media, we can now examine key questions about the behavioral and neural underpinnings of diverse and extensive real-world memories. Here, we tested the neural representations of episodic memory in a naturalistic setting, specifically focusing on the age, location, subjective memory strength, and emotional content of memories. We recruited 23 users of a video diary app (“1 Second Everyday”), who had recorded a total of 9,266 daily memory videos spanning up to 7 years prior to our study. During a 3T fMRI scan, participants viewed a set of 300 of their own memory videos intermixed with 300 videos from another individual. We identified key areas specifically engaged for one’s own memories versus another’s. Delving into the multidimensional nature of these memories, we find that their features are tightly interrelated, highlighting the need to consider these features in conjunction when conducting memory research. Importantly, when looking at the distinct contributions of these features, we find a topography of memory content extending across the medial parietal lobe with separate representations of a memory’s age, it’s strength, and the familiarity of the people and places involved.

## Introduction

Our episodic memories are dynamic and complex, filled with movement, emotions, meta-cognitive states, and multisensory information for the people, places, and events across the decades of our lives. However, memory research conducted in laboratories often cannot capture the richness of these real-world memories, instead prioritizing well-controlled stimuli at a short time scale. Other studies using autobiographic memories examine longer time scales, but usually with a constricted set of memories dichotomized along a single dimension such as time. How can we capture the rich complexity of real-world memories in an experimental setting, through the lens of the thousands of memories people naturally record in their daily lives?

For a majority of autobiographic memory studies, participants verbally self-report memories, bring in photographs of important events from an album, or freely recall a memory based on a vague cue like “beach” (see Cabeza & St. Jacques, 2007 for a review of methods). However, such methods assume an accurate report from the participant, and often focus on a handful of particularly salient events that may trigger unique processing and representations in the brain, given their high vividness (Sreekumar et al., 2018). To capture more naturalistic, daily events, some studies have employed wearable cameras that capture photographs at regular time intervals (Nielson et al., 2015; Rissman et al., 2016), but these studies often suffer from an opposing issue, where many photos are unmemorable to participants, and these studies can only measure the brief time span a participant is willing to wear a camera (less than a month). Due to these constrained methodologies, autobiographical memory studies have largely been limited to relatively small numbers of stimuli, defined by coarse condition contrasts such as autobiographic versus laboratory stimuli, face versus scene memories, or remote versus recent memories.

However, with the explosive growth of social media, high quality mobile cameras, and expansive cloud storage, people are recording their experiences more often than ever. One popular mobile application for capturing and sharing memories is “1 Second Everyday” (1SE) — an app developed in 2013 in which users record a 1-second video every day of their lives as a video diary. While one second is brief, these videos are information-dense, with dynamic visual content that serves as highly salient cues to specific memories. A single second allows one to easily capture and store thousands of these memory cues— a decade of memories lasts about one hour, and a lifetime about 8-9 hours. Because these memory cues are recorded every day, they capture the naturalistic events of one’s life—not just the key events that would be recalled from a photo album or cued memories. With over 1.5 million users, there is also a large body of individuals who have been using this app to capture their lives for years. We leverage this rich pool of participants with thousands of recorded videos to examine the neural representations of memories across their multifaceted components.

Prior research has identified several key neural substrates related to autobiographical memory representations: in particular, the hippocampus and the parietal cortex. The hippocampus is actively engaged in the encoding and retrieval of autobiographical memories, and shows increased signal for recall of memories that are more vivid (Shrager et al., 2008; Geib et al., 2017) and more recent (Bonnici et al., 2012; Nielson et al., 2015). This recency effect has received close examination, to adjudicate between competing theories of memory consolidation which posit that the hippocampus is either involved in the retrieval of recent memories (standard trace theory: Alvarez & Squire, 1994), or in episodic memory retrieval regardless of memory age (multiple trace theory: Nadel & Moscovitch, 1997). These accounts are difficult to test because of the interrelated nature of the properties that form memory—for example, memory strength and recall vividness can often be confounded with recency (Yonelinas et al., 2019), and thus must be taken into consideration (Gilmore et al., 2021). In addition to the hippocampus, attention in the field has also focused around both lateral and medial parietal regions (Wagner et al., 2005; Cavanna & Trimble, 2006; Gilmore et al., 2015).

Previously thought to be primarily engaged in working memory or the direction of attention to memory (Wagner et al., 2005), recent accounts have revealed signatures for long-term memory of familiar people and places in medial parietal cortex (Maddock et al., 2001; Silson et al., 2019a; Woolnough et al., 2020; Steel et al., 2021). Rare patients with lateral parietal lesions (Berryhill et al., 2007; Philippi et al., 2015) and medial parietal lesions (Valenstein et al., 1987) also show a marked loss in the retrieval of autobiographical memory details, and such deficits can be triggered in healthy individuals by transcranial magnetic stimulation to the lateral parietal cortex (Bonnici et al., 2018). Thus, while the hippocampus may be sensitive to the age or strength of a memory, it appears that the parietal cortex is sensitive to the content of a memory.

Now, armed with a rich, varied set of daily documented memories spanning a wide but finely sampled temporal range, we can examine key questions about neural representations of memory. Do we observe differences in hippocampal engagement between recent and remote memories, and how do such patterns relate to perceived memory strength? What fine-grained memory content can we decode from the parietal cortex? And more broadly, what representational information do we conjure from a memory during retrieval?

Here, we present a comprehensive characterization of the neural substrates for retrieving memories documented across hundreds or thousands of days, spanning as broad a range as seven years. We recruited users of the 1SE app with at least 6 months of recorded memories, and conducted a functional magnetic resonance imaging (fMRI) experiment in which participants watched their own recorded videos (Figure 1). Importantly, participants viewed their videos interleaved with an equal number of videos from a matched participant, to be able to isolate neural patterns for mnemonic representations rather than perceptual ones. From each participant, we collected rich behavioral measures for each video, including its age (its time of occurrence), location, memory strength, emotion, and content (i.e., how familiar were people and places shown in the videos?). These neural and behavioral data form one of the largest scale databases of autobiographic memory to date (https://osf.io/exb7m/?view_only=8d6939d3fd7c4819aa25aaa5391d10a3). We observe a high degree of correlations across our behavioral measures previously underreported in the field, suggesting that some prior findings could be accounted for by memory strength or even stimulus characteristics, rather than the memory property of interest. Furthermore, we uncover a topography within the medial parietal cortex that reflects memory content, age, and memory strength.

**Figure 1.**
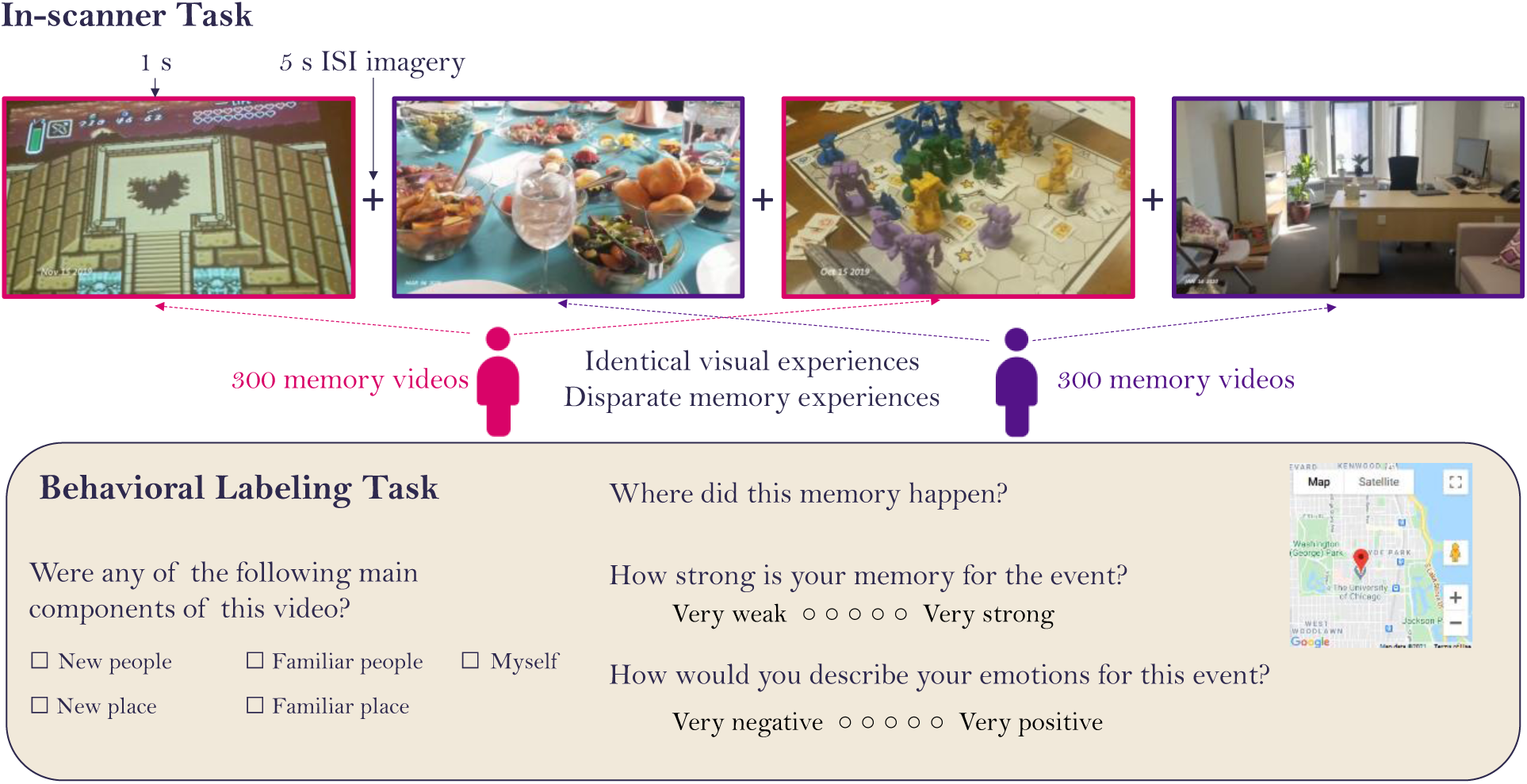
Experimental methods. For the in-scanner task, participants viewed a randomly intermixed sequence of 1-second videos, consisting of approximately 300 of their own memory videos and 300 memory videos from a paired participant. This participant pair saw the exact same videos in the same order, so their visual experiences were identical, but their memory experiences were non-overlapping, only recognizing their own videos and not recognizing the videos of the paired participant. After viewing a 1s video, participants had a 5s fixation period during which they were asked to imagine or recall the context surrounding the 1s video clip. Participants completed a behavioral labeling task after the scan, where they rated their own 300 videos on a series of questions, including the content of the video (people and places in the video), the video location (using GPS coordinates on a map), their memory strength, and emotions for the event.

## Results

Twenty-three users of the 1SE app participated in our study. Each participant had on average 762 (SD=536.2, min=175, max=2210) recorded videos to choose from: 11 participants with 6 months to 1 year of videos, 6 participants with 1-2 years of videos, and 6 participants with 2-7 years of videos. For each participant, up to 300 of their videos were pseudo-randomly sampled for use in the study, so that their entire time span was evenly sampled in each of the 10 fMRI runs. Nine participants returned for a second session (tested with 300 different videos) at least 6 months after the first session. All analyses reported here are conducted at the level of experimental session (32 samples), with first and second sessions of a given participant treated as independent sessions. Results were qualitatively similar if only a single session was included from each participant (Supplemental Figure 4).

We conducted a series of analyses on the behavioral and neuroimaging data, which we describe in detail below. First, we examined the distribution of the behavioral characteristics of the memories, revealing correlations across several measures. Second, we identified key regions in the brain specifically sensitive to the retrieval of one’s own autobiographic memories. Focusing on the medial temporal lobe, we found discriminable information about memory strength and emotion, but not age or spatial location. We also found that memory strength for a video could be predicted by its visual features. Third, we examined the unique contributions of these different memory features (memory strength, age, location, emotion, and content) across the brain, and found that the medial parietal cortex showed a bilateral topography of distinct representations of memory age, memory strength, people familiarity, and place familiarity. All brain data from the 32 samples and measures from the 9,266 total videos will be made publicly available on a repository on the Open Science Framework (https://osf.io/exb7m/?view_only=8d6939d3fd7c4819aa25aaa5391d10a3).

### Memories show rich variation and interrelatedness of features

We assessed and compared the behavioral features of these memories recorded “in the wild” to determine what influences memory strength and emotion (Figure 2). Many studies have focused on singular aspects of a memory, without considering that multiple aspects of a memory may be interrelated. For each of a participant’s approximately 300 videos that they viewed in the scanner, participants made ratings on several aspects of the associated memory (see *Methods*). Participants indicated the content of the videos, by rating the familiarity of people and places in their videos (i.e., was this a video of meeting someone for the first time?). We also asked participants to label each video for: 1) its specific location (geocoordinates), 2) the strength of the memory of the event, and 3) the emotionality of the captured event (valence: positive vs. negative, and strength: strong vs. weak). We quantified the distance of a video as the geodesic distance between the video location and the study site where the participant was scanned. The study site was located in the Washington, DC metropolitan area (Bethesda, MD), and all participants lived in the DC area. We also obtained a measure of the age of a video, from the time the video was recorded in the app.

**Figure 2.**
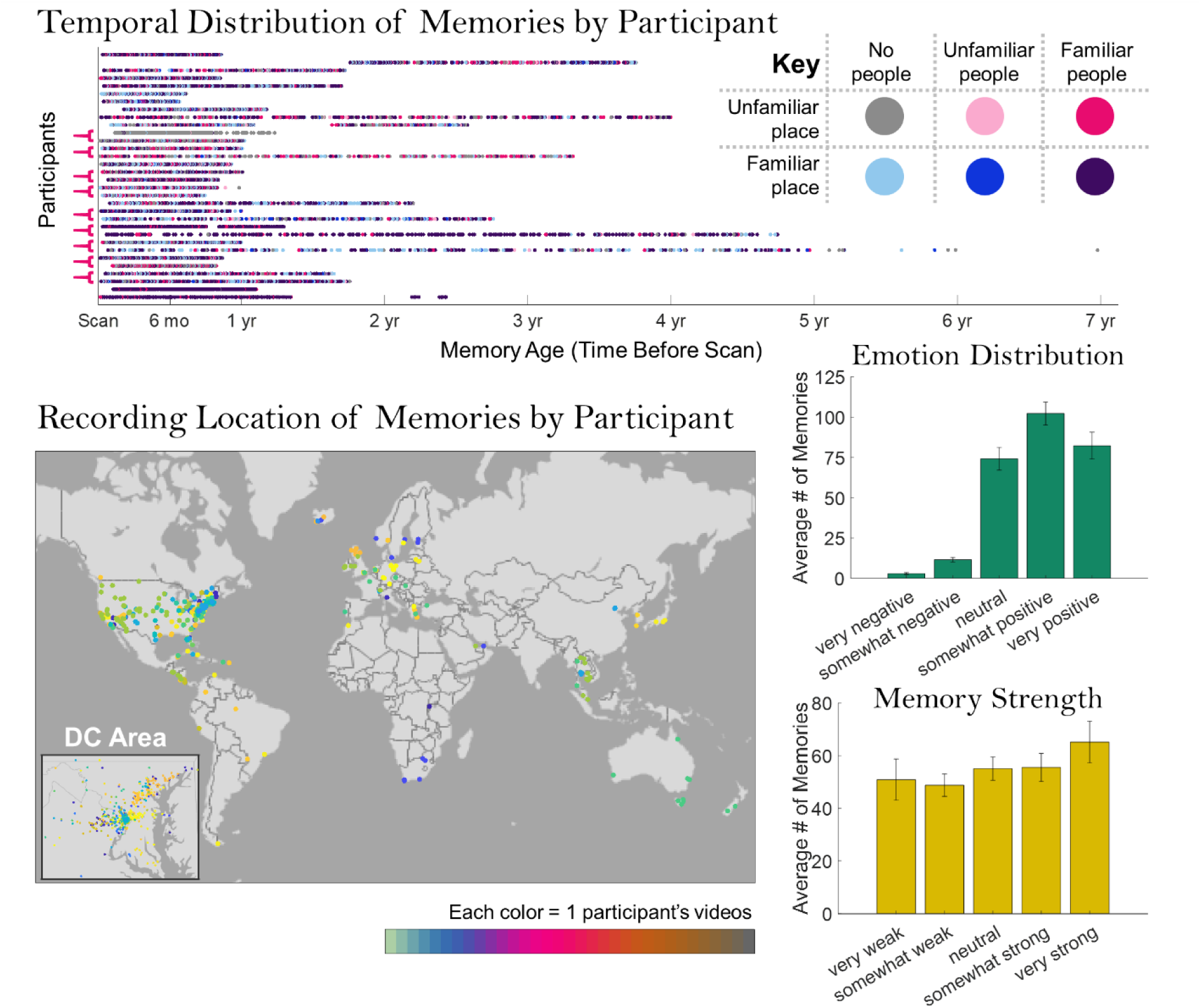
The diversity of memories in the study. These plots show the distributions of the ∼300 memories per participant used in the current study along a range of metrics. The temporal distribution shows a scatterplot of the age of each of the participants’ memories. Each horizontal row represents each of the 32 experimental samples, and samples from the same participant (N=9) are indicated with brackets at the left. While many memories occurred within one year of the experimental scan, several participants had memories extending more than a year prior, and some up to seven years prior to the experiment. Dots are color-coded based on memory content (whether there are familiar / unfamiliar people in the video, and whether it occurs in a familiar place). The content types are diverse across participants, with some recording a majority of videos with familiar people in familiar places, others recording a majority of videos with novel people, places, or both, and even others recording a mix of all content types. The spatial distribution shows the locations of all videos, with each dot representing a video and each color representing a participant. Videos were in diverse locations, with many across the world, even more across the United States, and a large number concentrated in the Washington DC area (where the study was conducted). Looking at a distribution of the emotional ratings for the memories, the documented videos in general tended to be neutral or positive (rather than negative). Memory strength ratings were evenly distributed from very weak to very strong. Error bars indicate the standard error of the mean across participants.

We first examined the spread of emotion and memory strength ratings for the videos. The videos generally skewed from neutral to positive in emotional valence (on average, 1.1% were very negative, 4.1% were somewhat negative, 27.5% were emotionally neutral, 37.8% were somewhat positive, and 29.5% were very positive). This positive bias may reflect an interesting aspect of memory; perhaps when documenting one’s life, app users prefer to select more positive videos to represent a day’s memories. In contrast, there was a relatively uniform distribution of memory strength ratings for these recorded videos (on a scale of 1 = very weak to 5 = very strong, 18.0% received a 1, 17.4% received a 2, 20.9% received a 3, 20.2% received a 4, and 23.5% received a 5).

Next, we investigated the relationship between the content of the videos (the presence of familiar people and places) and the other memory properties. On average, 74.3% of videos (*SD* = 21.6%) had people in them, with 93.2% (*SD* = 6.4%) rated as familiar people (Supplemental Figure 1a). Videos that contained new people were significantly better remembered (i.e., had a higher rated memory strength) than those with familiar people (*t*(28)=3.07, *p*=0.005). However, there was no significant difference in the emotionality of a video based on the familiarity of people in the video (*p*=0.754). Participants also sometimes recorded themselves in their videos (as “selfies”). Participants reported higher memory strength for videos with new people than those with themselves (*t*(28)=2.33, *p*=0.027), and there was no difference in memory for videos with themselves or familiar people (*p*=0.839). Videos with selfies did not show differences in emotion from videos with familiar (*p*=0.144) or new people (*p*=0.744).

In terms of the places shown in these videos, a majority of videos were recorded in familiar places (*M*=80.0%, *SD*=12.0%, MIN=87, MAX=286). Videos that occurred in new places had significantly higher memory strength than those in familiar places (*t*(31)=9.89, *p*=4.13 × 10^-11^). New place videos were also significantly more positive (*t*(31)=5.03, *p*=1.96 × 10^-5^) and in farther locations from the study location (*t*(31)=5.29, *p*=9.45 × 10^-6^), perhaps reflecting positive memories of travel and vacations. We also categorized memories that occurred in unique locations (locations with only a single memory, such as a vacation spot), and common locations (the most common location across all memories for a participant, such as one’s home; Supplemental Figure 1b). Videos that occurred in unique locations had significantly higher memory strength than those that happened in common locations (*t*(31)=12.58, *p*=1.02 × 10^-13^), and were rated with more positive emotions (*t*(31)=6.07, *p*=9.97 × 10^-7^), again perhaps reflecting a boost to unique vacation memories.

Finally, we investigated the interrelationships of the qualities of the memories, their emotionality, strength, age, and distance. There was no significant difference in memory strength between negative and positive videos (including only participants with at least 5 negative and 5 positive videos: *t*(26)=1.32, *p*=0.200). Clear relationships emerged in terms of memory strength and emotion as compared to a memory’s age and distance (Figure 3). In terms of memory age, more recent videos were rated as being better remembered (Mean Spearman’s *ρ=*0.16, *t*(31)=4.54, *p*=7.96 × 10^-5^) and had more positive emotions (Mean *ρ*=0.07, *t*(31)=3.71, *p*=8.12 × 10^-4^). In terms of distance, farther away videos were better remembered (Mean *ρ*=0.19, *t*(31)=8.08, *p*=3.95 × 10^-9^) and more positively remembered (Mean *ρ*=0.19, *t*(31)=8.39, *p*=1.78 × 10^-9^), echoing the behavioral findings for novel places.

**Figure 3.**
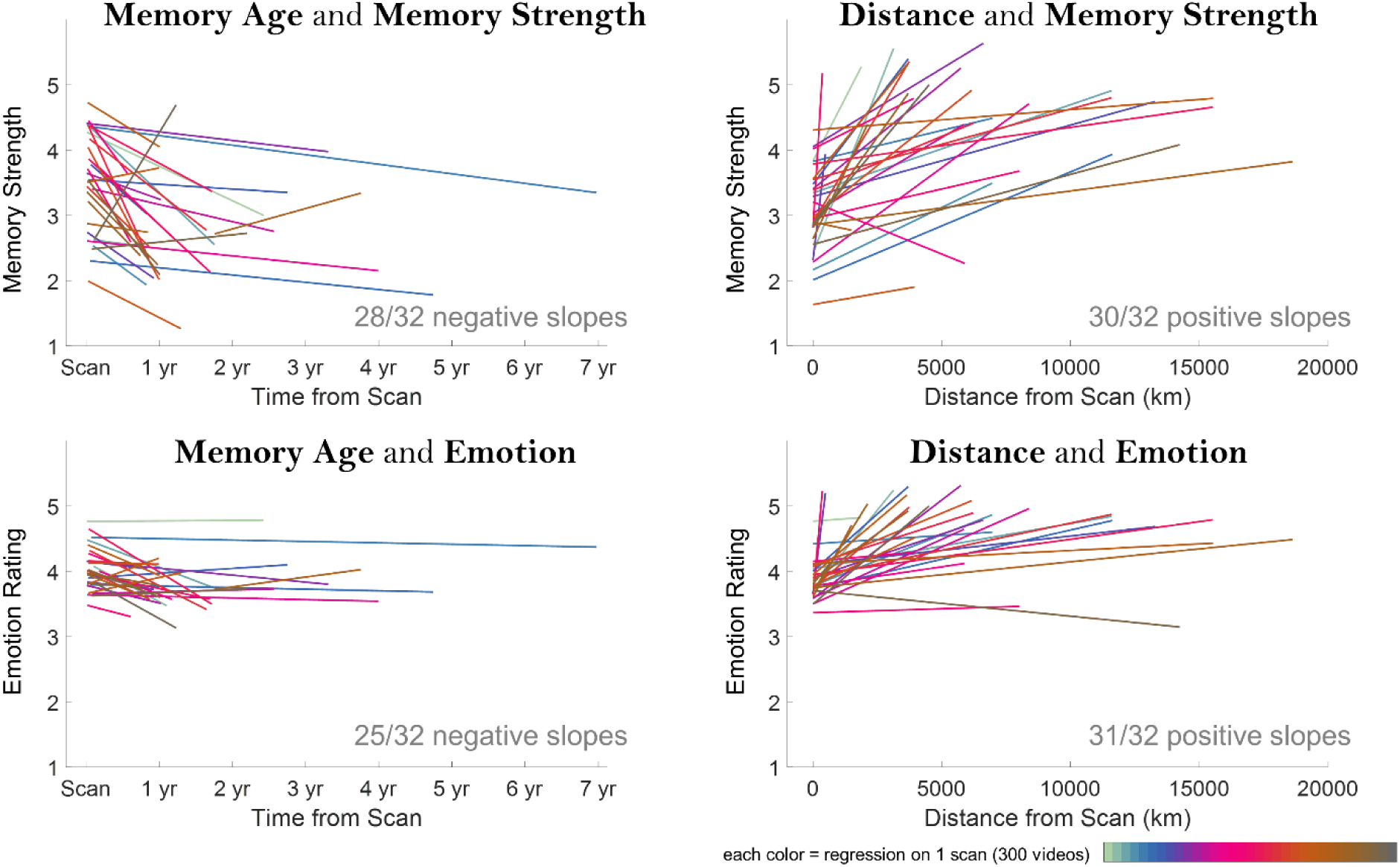
Relationships across memory qualities. Each plot shows regression fit lines (intercept and slope) for all 32 samples for four comparisons: (top left) memory age and strength, (top right) memory distance and strength, (bottom left) memory age and emotion rating, and (bottom right) memory distance and emotion rating. These trends are shown for illustrative purposes, and are statistically confirmed by Spearman rank correlations in the main text that make no assumption about linearity. Memory strength ratings ranged from 1 (very weak) to 5 (very strong), while emotion ratings ranged from 1 (very negative) to 5 (very positive). In 28 out of 32 participant samples, age had a negative relationship with memory strength (i.e., older memories were less strongly remembered). In 30 out of 32 samples, distance had a positive relationship with memory strength (i.e., farther memories were more strongly remembered). In 25 out of 32 samples, age had a negative relationship with emotion (i.e., older memories had weaker emotions). In 31 out of 32 samples, distance had a positive relationship with emotion (i.e., farther memories had stronger positive emotions). Note that the emotion regression lines tend to fall in the upper half of the charts due to the overwhelmingly neutral and positive memories reported by participants (ratings of 3-5).

These results emphasize that the attributes of memories are strongly interdependent: one cannot study the effects of memory age or location in isolation without also considering memory strength or emotion. Similarly, the specific content of the memory (who is there, where is it happening) also may impact how that memory is represented. In this sample, the most memorable events appear to be recent, positive events with new people in far-off new locations. The most forgettable ones are those that occurred long ago, with highly familiar people and places.

### Representations in the brain for one’s own memories

With these highly interconnected features that are inherent to a memory, how do they interact to form a neural representation of a memory? To address this question, we first tested whether we could identify neural representations specific to an individual’s own memories. In the MRI scanner, when participants viewed one of their own 1-second videos, they were instructed to recall the events surrounding the video in as much detail as possible. When they viewed a 1-second video from a paired participant, they were instructed to instead imagine the events surrounding the video. Thus, neural signatures for both types of stimuli would reflect visually and semantically rich representations and constructive, imagery-based processes. Both participants in a pair in fact saw the exact same visual stimuli (the same videos in the same order), so the only differences between a pair should be the individual-specific mnemonic associations with the videos. Thus, we examined where we could observe distinguishable traces in the brain for viewing one’s own memory videos versus another’s memory videos.

A group-level contrast of viewing one’s own videos versus the other’s videos (Figure 4; unthresholded maps in Supplemental Figure 2) revealed significant, bilateral clusters of activation in the ventromedial prefrontal cortex (vmPFC), medial parietal cortex (mPC, encompassing areas such as the retrosplenial cortex, posterior cingulate cortex, and precuneus), lateral parietal cortex, the medial temporal lobe (MTL), and hippocampus. These regions highly overlap with regions previously identified for autobiographical memory processes, including many regions often summarized as the default mode network (DMN; Spreng et al., 2008; Philippi et al., 2015). Importantly, the observation of mPC activity in our task supports prior findings of mPC involvement in autobiographical memory (Cavanna & Trimble, 2006), and shows significant overlap with mPC regions implicated in sensitivity to familiar people and familiar places (Supplemental Figure 3; Silson et al., 2019a). Similarly, these MTL and hippocampal effects support prior work showing the involvement of these regions in autobiographical memory strength or age (Shrager et al., 2008; Bonnici et al., 2012; Geib et al., 2017). Importantly, we do not observe significant effects in parts of the occipital lobe corresponding to early visual cortex (V1-V3), suggesting that indeed there were no differences in the visual experiences while watching the videos between paired participants. Significant regions for the opposite contrast of higher activity for viewing another’s videos over one’s own videos do emerge in the insula and in lateral parietal regions (blue areas in Figure 4). These regions could potentially reflect a novelty effect, an effect of increased imagining for others’ events (Addis et al., 2007; Gilmore et al., 2016), or a deactivation effect related to rerouting of resources during episodic memory (White et al., 2013; Chai et al., 2014). As a whole, clusters observed in this contrast replicate across scanning sessions for the nine participants who participated in two sessions (Supplemental Figure 4). In sum, these clear differences between viewing one’s own videos and those of another indicate that these stimuli elicit strong signatures related to memory for the video’s events, rather than only a perceptual representation.

**Figure 4.**
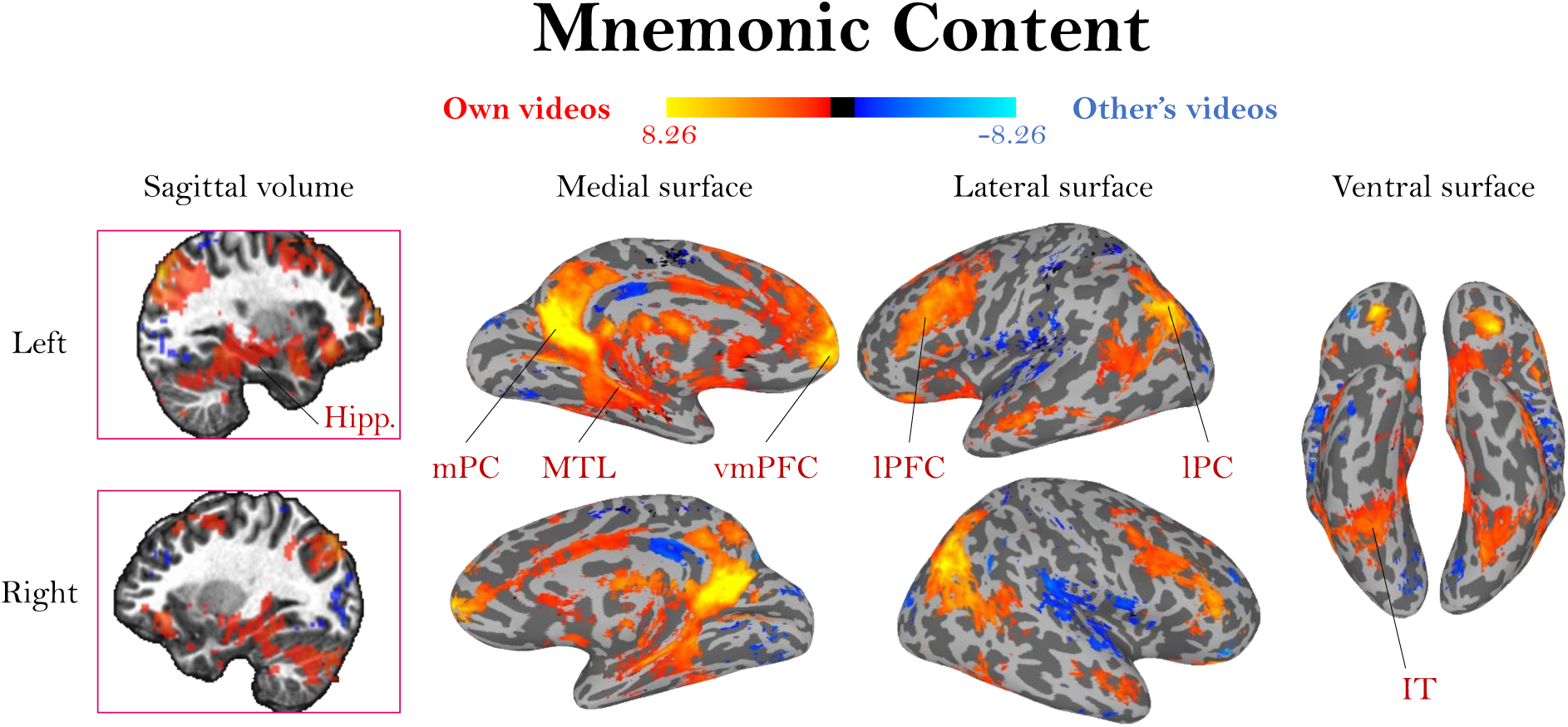
Activation differences based on video mnemonic content. A whole-brain group activation map (N=32) for viewing one’s own videos (red/yellow) versus viewing another person’s videos (blue), FDR-corrected, *q* < 0.01. The colormap represents the range of beta values. Because participant pairs had identical visual content, these patterns should solely represent activation related to memory for the event. Activation for viewing one’s own videos coincides with regions frequently observed in autobiographical memory studies, including hippocampus (Hipp.) and medial parietal cortex (mPC).

### Mnemonic representations of time and space in the medial temporal lobe

Next, we investigated targeted regions revealed in prior work to maintain representations of memory age and location—namely the hippocampus (Bonnici et al., 2012; Eichenbaum, 2013; Nielson et al., 2015) and other surrounding medial temporal lobe (MTL) regions: the parahippocampal cortex (PHC), entorhinal cortex (ERC), and amygdala (see Supplemental Tables 1-3 for betas and results split by hemisphere).

For each region of interest, we ran regressions to test the relationship of voxel signal and our measures of memory: age, distance, memory strength, and emotional valence. First, we examined each of these factors in separate models, in order to replicate prior work that has focused on singular properties like memory age or strength, before testing in a combined model. The regression slopes for all participants were then compared to a null hypothesis of zero slope. All MTL regions showed a significant effect of age, where more recent memories had higher signal (Hippocampus: *p*=0.009, Cohen’s *d*=0.50; Amygdala: *p*=0.013, *d*=0.47; ERC: *p*=0.010, *d*=0.48; PHC: *p*=0.005, *d*=0.54; all FDR *q*<0.05). All regions also showed a significant effect of memory strength, where stronger memories had higher signal (Hippocampus: *p*=4.46 × 10^-6^, *d*=0.98; Amygdala: *p*=1.09 × 10^-5^, *d*=0.93; ERC: *p*=6.37 × 10^-6^, *d*=0.96; PHC: *p*=8.10 × 10^-5^, *d*=0.80; all FDR *q*<0.05). These regions also showed an effect of emotion, where more strongly positive memories had higher signal (Hippocampus: *p*=7.27 × 10^-5^, *d*=0.81; Amygdala: *p*=2.18 × 10^-5^, *d*=0.88; ERC: *p*=4.52 × 10^-5^, *d*=0.84; PHC: *p*=0.008, *d*=0.50, all FDR *q*<0.05). There were no regions with bilateral significant effects related to the distance of a memory (all *p*>0.05). In sum, the MTL shows sensitivity to a memory’s age, strength, and emotion when these properties are tested separately.

Given our behavioral findings that these different memory properties are in fact highly correlated, we next examined how these different factors played a role in predicting MTL signals when tested in a combined model. With a combined model (Figure 5a), across all regions the significant effect of age disappeared (all *p*>0.10), and there continued to be no significant effect of distance (all *p*>0.15). In contrast, significant effects of memory strength remained in all regions (Hippocampus: *p*=1.57 × 10^-4^, *d*=0.76; Amygdala: *p=*6.12 × 10^-4^, *d*=0.67; ERC: *p*=0.003, *d*=0.56; PHC: *p*=0.014, *d*=0.46, all FDR *q*<0.05). These findings corroborate a large body of work showing memory strength effects in the hippocampus (e.g., Shrager et al., 2008; Geib et al., 2017). Significant effects of emotional strength also remained in all regions except the PHC (Hippocampus: *p*=0.009, *d*=0.50; Amygdala: *p*=4.59 × 10^-4^, *d*=0.69; ERC: *p*=0.003, *d*=0.56; all FDR *q*<0.05; PHC: *p*=0.218). These findings replicate prior work showing representations of the emotional content of a memory in the amygdala, hippocampus, and other MTL regions (Piefke et al., 2003; Daselaar et al., 2008). However, the disappearance of an effect of memory age when tested in this combined model suggests that memory age effects in the MTL could be partially accounted for by confounding effects of memory strength or emotion.

**Figure 5.**
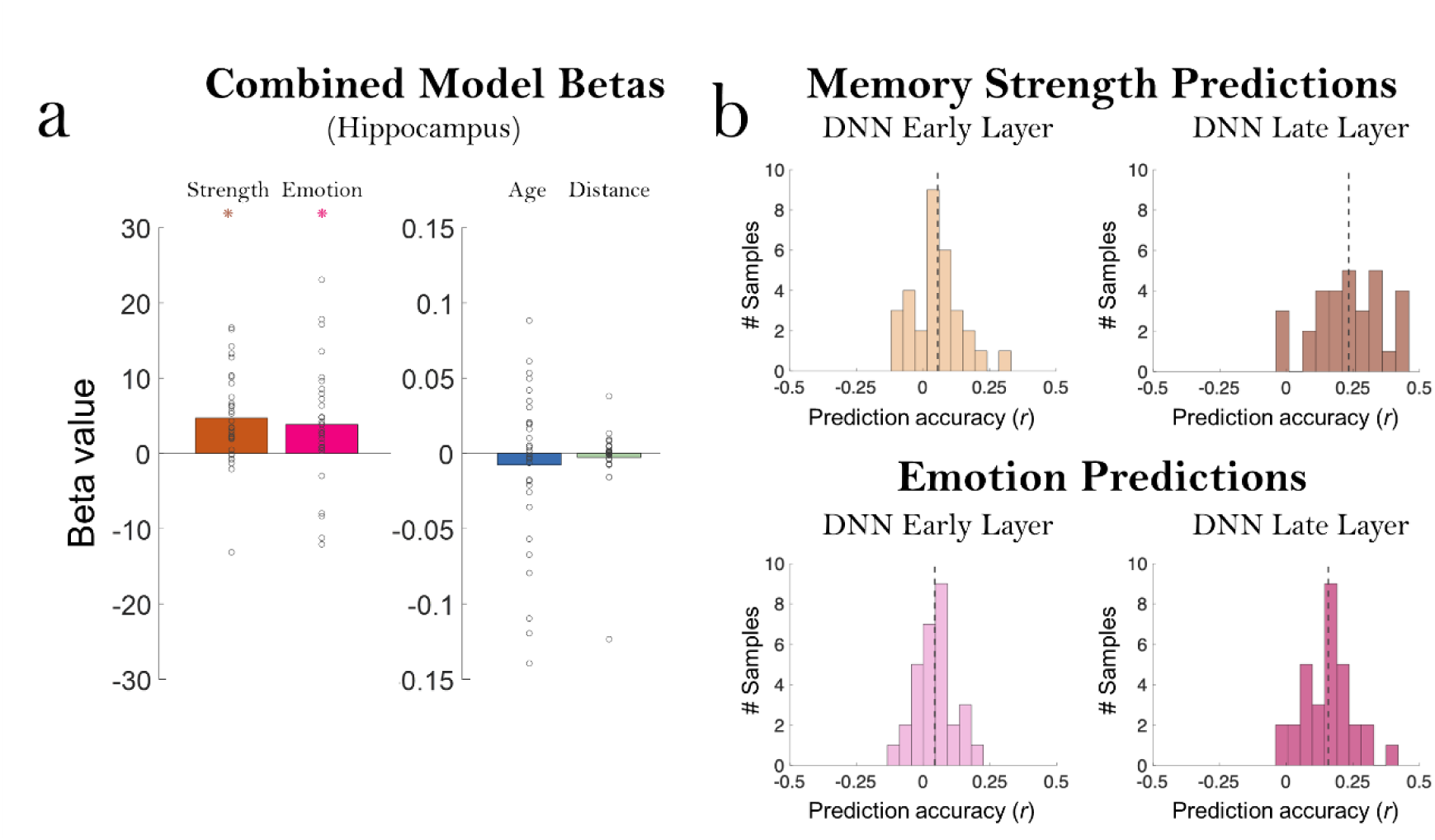
Significant effects of memory and emotion in the hippocampus and DNN predictions. a) Betas for all participants in the combined model predicting hippocampal activation from the four factors of memory strength, emotion rating, age (time from scan), and distance from scan site. Each point indicates one of the 32 samples, and the bar indicates the average beta value across participants. Note the much more constrained y-axis needed to display the age and distance factors (beta range of -0.15 to 0.15) versus the memory and emotion factors (beta range of -30 to 30). While memory strength and emotion show a significant positive relationship to hippocampal activation (*, *p*<0.01), age and distance do not (*p*>0.10). b) Prediction accuracy of the VGG-16 Deep Learning Neural Network (DNN) layer activations on the middle video frame, for predicting memory strength and emotion ratings. Each histogram shows the distribution of prediction accuracy (Pearson correlation *r* of predictions with true values of memory strength / emotion) of the 32 participant samples. The dashed gray line indicates the mean prediction accuracy across all participants. For memory strength, both early (layer 2) and late (layer 20) layers were significantly able to predict memory strength from the middle frame of the videos alone (top). Early and late layers were also significantly able to predict emotion ratings from the middle video frame (bottom).

We also tested the ability to predict MTL signals with a combined multiple regression model using a representational similarity analysis (RSA) framework (Kriegeskorte et al., 2008; Nielson et al., 2015). Instead of predicting mean activation for a memory video in an ROI based on properties of that memory, RSA looks at predicting pairwise pattern differences from pairwise behavioral differences (see *Methods*). For example, one prediction might be that two memories that are very different in emotional content would be very different in neural signal. When testing a combined model including representational dissimilarity matrices (RDMs) for memory strength, age, distance, and emotion, we only find a significant contribution of memory strength (Hippocampus: *p*=1.16 × 10^-4^, *d*=1.14; Amygdala: *p*=3.08 × 10^-4^, *d*=0.92; ERC: p=3.58 × 10^-4^, *d*=1.15; PHC: *p*=0.013, *d*=0.57; all FDR *q*<0.05). Memory age, distance, and emotion showed no significant relationship (all *p*>0.05). Overall, these results suggest that effects of memory age or distance observed in the hippocampus and MTL may be largely captured by memory strength and emotion of the memories, as more recent memories and more spatially distant memories tend to also be the best remembered and the most positively rated.

As a control analysis, we can also examine whether any aspects of a memory can be detected when a participant is viewing the videos of another person for which they have no memory. This can reveal whether there may be a contribution of visual features to these features of the memory that account for MTL activation. We tested this by examining a combined model for predicting the brain activity from a participant *A* when viewing the videos of participant *B*, with predictors for the memory strength, emotion, age, and geodesic distance as indicated by participant *B*. One should expect no ability to predict brain activity in this analysis, because participant *A* should have no knowledge of this information specific to participant *B* (such as the strength of participant *B*’s memory). However, surprisingly, in this combined model, there is a weak but significant relationship in some MTL regions between the voxel patterns of participant *A* watching these videos and participant *B*’s rated memory strength, in spite of no prior experience with these videos (Hippocampus: *p*=0.030, *d*=0.41; Amygdala: *p*=0.050, *d*=0.55; PHC: *p*=0.047, *d*=1.39). No other memory features were decodable. This suggests that memory strength is partially reflected in the stimulus itself; there may be certain visual features that correlate with memory strength, such as an intrinsic memorability of these stimuli (Bainbridge, 2019). This could also suggest that participants are vividly imagining or simulating the events surrounding these other-participant videos, which could engage the hippocampus (Martin et al., 2011).

Given these results, we tested whether objectively measured visual features from the videos could be used to predict subjective memory properties like memory strength or emotion. We utilized the image classification deep neural network (DNN) VGG-16 (Simonyan & Zisserman, 2014) to quantify the visual information present at the middle frame of each 1- second video (chosen to be representative of the whole video). While such object classification DNNs are originally designed to classify the visual features of images, they can also be successfully utilized to predict the intrinsic memorability of images (Khosla et al., 2015; Needell & Bainbridge, 2022). To quantify these videos, we used a cross-validated support vector regression (hold-out: 80% training, 20% testing, 25 iterations) to predict human memory strength ratings from features in an early layer (layer 2) thought to represent low-level visual information, and a late layer (layer 20) thought to represent high-level or more conceptual information (Figure 5b). Early layer features were significantly predictive of the participant’s ratings of memory strength (Wilcoxon sign rank test: Z=2.90, *p*=0.004), as were late layer features (Z=4.74, *p*=2.11 × 10^-6^), and late layer features were more predictive than early layer features (Z=4.45, *p*=8.47 × 10^-6^). This DNN was also predictive of emotion ratings at both early (Z=2.91, *p*=0.004) and late layers (Z=4.84, *p*=1.30 × 10^-6^), with significantly better predictions by the late versus early layer (Z=4.64, *p*=3.52 × 10^-6^). We also examined the role of motion in the videos, as measured by optical flow across the first, middle, and last frames (see *Methods*).

Motion was not predictive of memory strength ratings (Z=0.98, *p*=0.327), but was predictive of the emotion rating of a video (Z=2.84, *p*=0.005) such that on average videos with more motion were rated as having stronger positive emotions. These results provide a possible explanation for why memory strength can be decoded from the brain of a participant who hasn’t experienced these memories; some 1-second videos or even some memories are more vivid and visually striking and result in higher memory strength and emotional content, and this can be detected by a computational vision model.

In sum, these results emphasize the importance of considering the multifaceted features of a memory: its strength, emotions, location, age, and even its sensory features. In fact, some memories or videos of memories that tend to be remembered more strongly may also tend to be more visually striking, and brain activity could reflect some of these differences in visual features. Considering these various aspects of memory may help avoid confounds that could explain prior findings, such as memory strength effects that could account for hippocampal signatures of autobiographical memory age and distance.

### Memory representations across the whole brain

Given the relatively weak effects of memory age and distance in the MTL, we next expanded our combined model to an exploratory whole-brain searchlight to identify whether such effects may occur elsewhere in the brain. In a targeted ROI analysis, we then tested the content specificity of the regions that emerged (see section below). First, using a liberal threshold (*p*<0.01, uncorrected), we identified voxels with significant slopes in a multiple regression predicting searchlight voxel values from memory distance, age, strength, and emotion, allowing us to examine the contributions of each factor (Figure 6; alternate views in Supplemental Figure 5; views at a stringent threshold in Supplemental Figure 6). Even at this liberal threshold, no regions emerged with a significant effect for a memory’s distance (although some temporal lobe regions do emerge if testing only with videos within a constrained range of 50km, Supplemental Figure 7). Clusters of voxels did emerge with sensitivity to the age of the memory. Specifically, bilateral regions in the medial parietal cortex showed higher signal for more remote memories. An area around the left temporal parietal junction and secondary somatosensory cortex, and an area in the left anterior temporal gyrus showed higher signal for recent memories. The hippocampus did not emerge as showing a specific effect of memory age. However, its activity did reflect memory strength, along with many cortical regions including medial parietal areas, anterior temporal lobe, medial prefrontal cortex, and lateral parietal regions (also present at FDR-corrected *q*<*0*.01, Supplemental Figure 6). Activity related to emotion also emerged in several regions, including the amygdala, temporal areas, and parietal regions (also present at *q*<0.01, Supplemental Figure 6).

**Figure 6.**
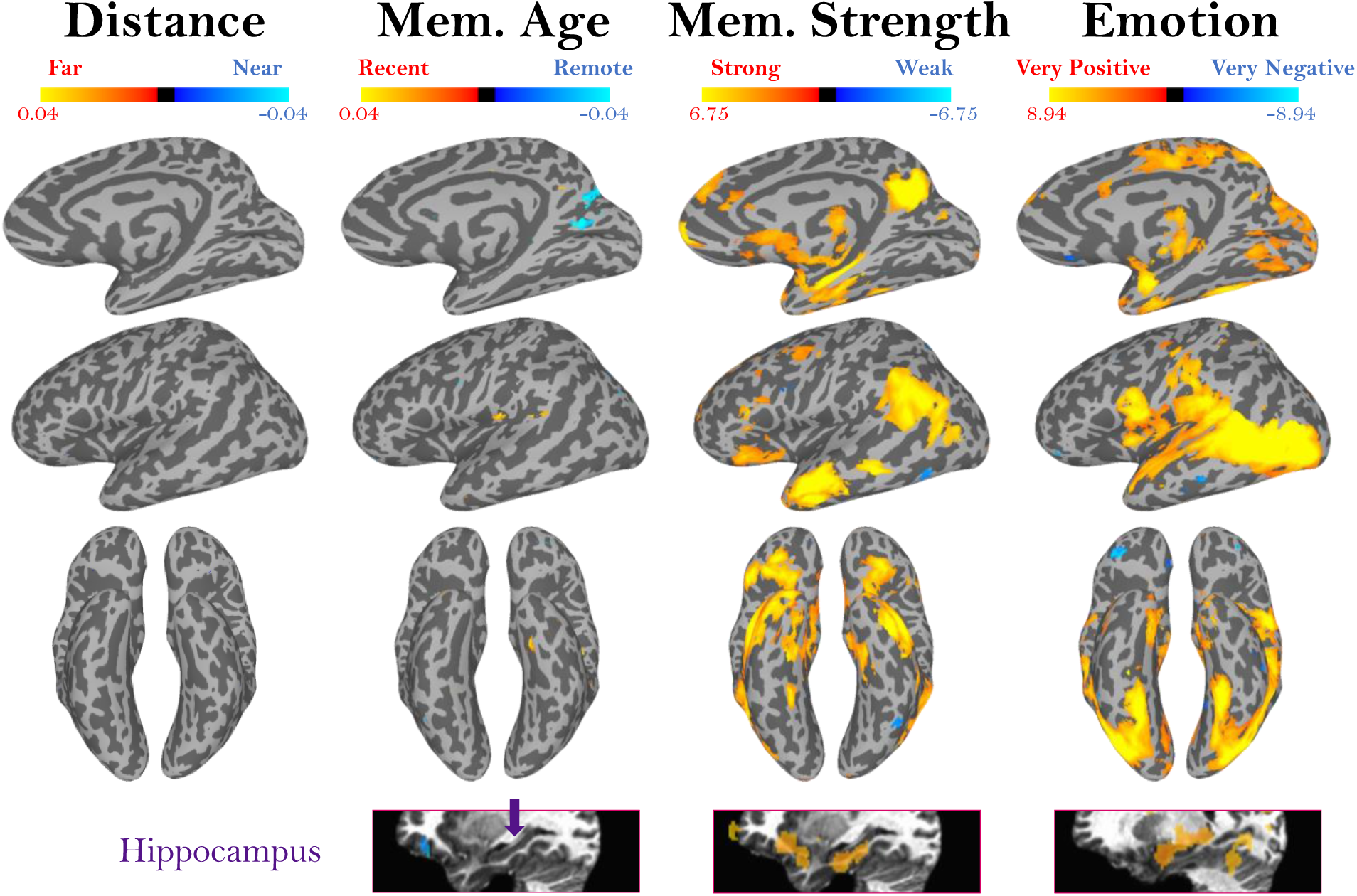
Representations of different memory content. Maps show whole-brain results from a multiple regression predicting voxel beta values from separate predictors for a memory’s distance, age, strength, and emotion. Activation represents the mean regressor slope (β) for each predictor, where significance was assessed by comparing the slope across all participants versus a null hypothesis slope of 0 (*p* < 0.01, uncorrected; more stringent threshold shown in Supplemental Figure 5). The colormaps represent the range of beta values for each predictor, after centering. Surface maps as well as a volume slice of the hippocampus (indicated in purple) are shown. These maps reveal voxels where the signal is significantly predicted by memory age, strength, and emotion. No regions emerged with sensitivity to memory distance. Alternate views can be seen in Supplemental Figure 5.

Interestingly, emotion is also the only memory feature reflected in activity in the visual cortex, possibly reflecting the relationship we observed between a video’s visual features and the emotion associated with a memory. These regions for memory age, strength, and emotion replicate when people and place familiarity are also included as predictors in the regression (Supplemental Figure 8).

This current analysis shows voxels with significantly higher and lower univariate activation related to *absolute* measures of the memory age, strength, and emotion for specific memories. A complementary measure is whether distances in neural patterns map onto behavioral distances between memories, reflecting sensitivity to *relative* differences between pairs of memories. To test this question, we ran a whole-brain RSA regression searchlight, to see how pairwise distances between memory videos in memory distance, age, strength, and emotion were predictive of pairwise distances in neural representation (see *Methods*). To calculate pairwise distances for memory age and distance, we modeled those values logarithmically, given prior work suggesting a logarithmic representation of time in behavior and the brain (Nielson et al., 2015; Singh et al., 2018). With this complementary analysis, we observe many similarities to the univariate analysis (Figure 7). Significant patterns for memory strength were observed in the medial parietal cortex, medial temporal lobe, and other temporal lobe regions. Patterns for emotion were observed in the temporal lobe and a more superior medial parietal region. Again, no significant regions emerged for memory distance.

**Figure 7.**
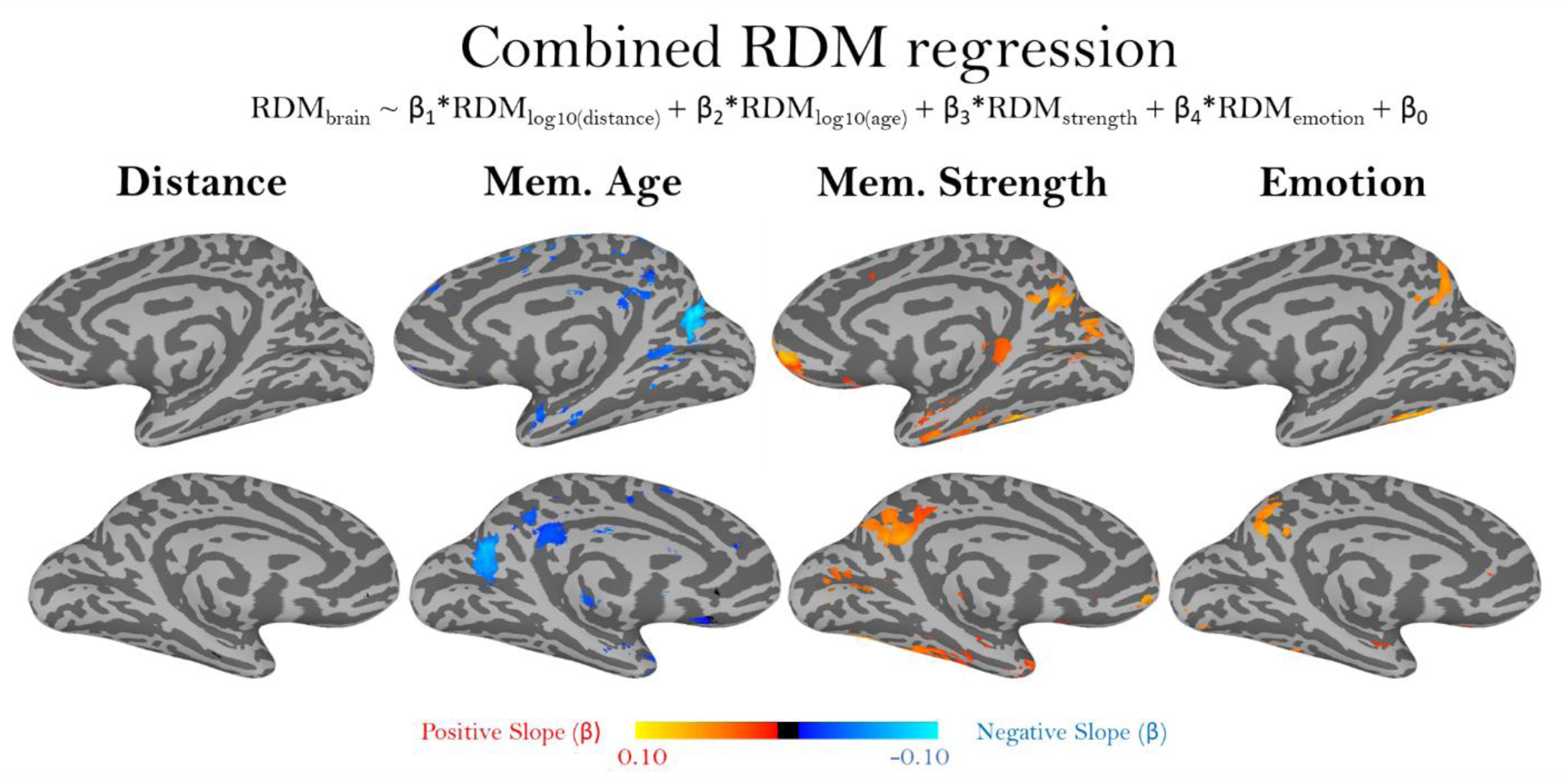
Representational similarity analysis of different memory content. Maps showing bilateral views of results from a multiple regression predicting the representational dissimilarity matrices (RDMs) of voxel beta values, from separate RDMs for a memory’s distance, age, strength, and emotion. Memory distance and age were logarithmically transformed before creating RDMs. Map values represent the similarity between the brain-based RDM and each predictor RDM, as the regressor slope (β). Significance was assessed by comparing the slope across all participants versus a null hypothesis slope of 0 (*p* < 0.01, uncorrected).

Finally, we again observe bilateral medial parietal patterns related to the age of a memory, as well as temporal lobe sensitivity. Interestingly, these patterns show a negative slope, indicating that memories that are closer together in time and/or memories that occurred longer ago are more neurally distinct.

### Medial parietal substrates of memory content and age

In both whole-brain analyses, bilateral clusters in the mPC emerged that were sensitive to the age of a memory and its strength. This temporal age region is of particular interest, given that we did not see sensitivity to memory age in other regions such as the hippocampus, even with a liberal threshold. Further, the mPC is thought to represent aspects of the content of a memory, particularly with distinct areas sensitive to the recall of familiar people and familiar places (Silson et al., 2019a; Woolnough et al., 2020). Our contrast of viewing one’s own versus another’s memories also showed selectivity in similar mPC areas (Figure 4). Thus, the mPC may be sensitive to different aspects of a memory’s content, and serves as a particularly interesting region to examine with our richly quantified memory set.

To test whether memory content effects are specific to the mPC, we examined the locations of the peak voxels for each type of memory information. Specifically, we identified the top 1000 voxels across the whole brain showing a significant effect for memory strength, memory age, and the familiarity of the people and places within the videos. These four contrasts revealed significant, symmetrical bilateral regions primarily within the mPC (Figure 8a, whole brain maps in Supplemental Figure 9, maps with the same colormap in Supplemental Figure 10). Importantly, these four contrasts had low voxel overlap with each other, suggesting separate clusters within the mPC for these different memory properties (age, strength, people familiarity, and place familiarity). The mPC did not contain any of the top voxels for emotion information (Supplemental Figure 11). Interestingly, the top significant regions for people familiarity showed a positive effect, while those for scene familiarity showed a negative effect (voxels for the opposite directions are shown in Supplemental Figure 12).

**Figure 8.**
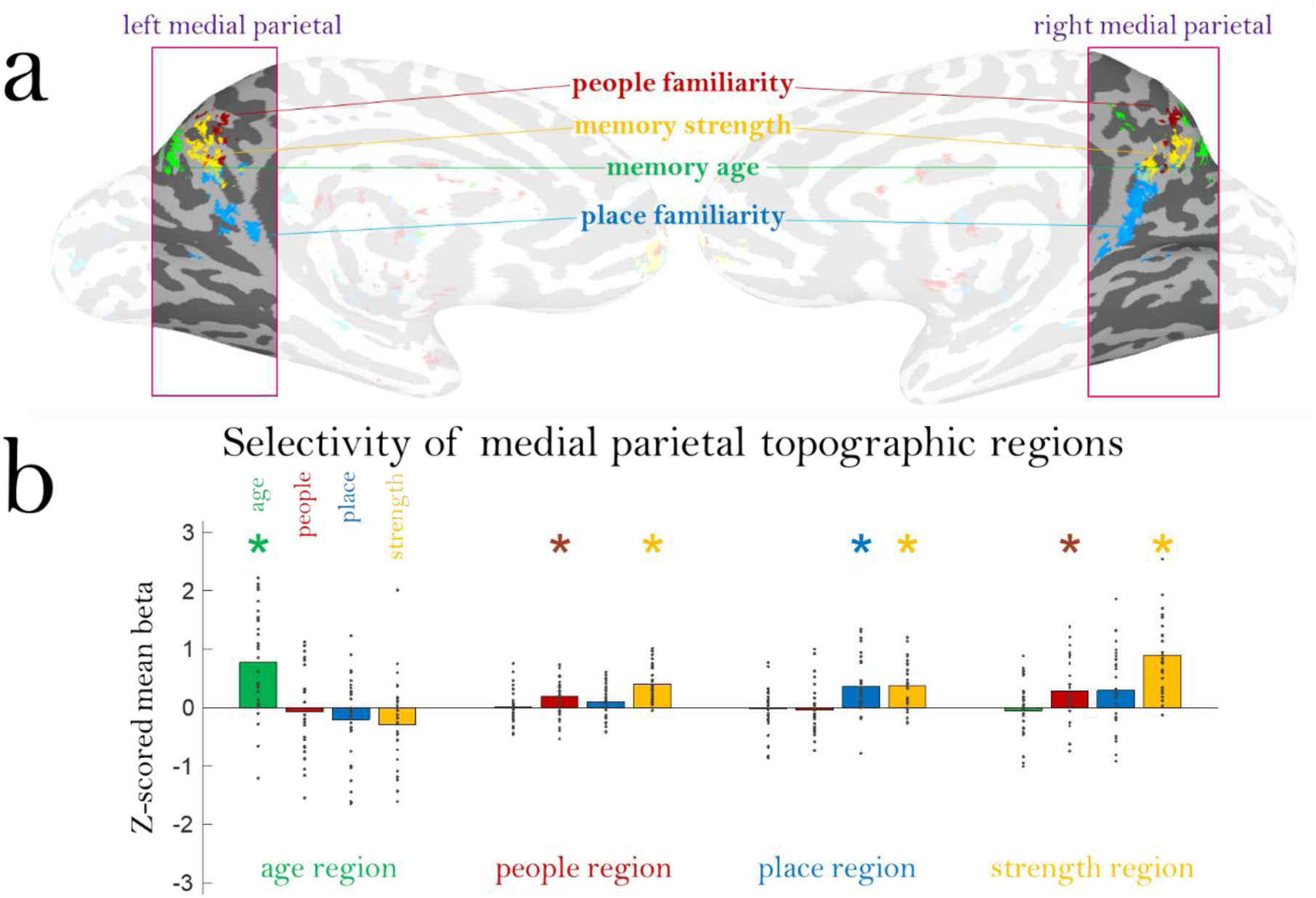
Distribution of memory content in the medial parietal cortex. a) The medial parietal cortex (mPC) contains a topographic map representing different types of memory information. Shown here are the top 1000 voxels with signal for four different types of information: memory age, people familiarity, place familiarity, and memory strength. Each map is shown at 50% transparency; if a voxel is shared across multiple content types, it will be colored by both maps (i.e., a “winner” is not determined in any given voxel). Note the low amount of overlap across content types and hemispheric symmetry of these maps. b) Results of a leave-one-out analysis of the mean beta for each content type in each of the topographic regions. Topographic regions were localized with all but one participant, and then the mean z-scored beta of the left-out participant is plotted (each dot). Bars show the mean beta across participants, and * indicate a significant difference in a two-tailed t-test versus 0 (FDR corrected across all comparisons, *q* < 0.05).

To confirm whether these mPC regions were specific to each factor, we tested selectivity of each region using a leave-one-participant-out approach. In other words, for each participant we localized each mPC region in all other participants, and then computed the mean z-scored beta value for the excluded participant for each factor (Figure 8b). We then compared the beta values across all participants versus 0 using two-tailed t-tests, FDR corrected *q*<0.05. This allowed us to see whether subregions within the mPC were topographically stable across participants (i.e., does participant *N* have discriminable information in a region defined by all other participants?), and whether these subregions were sensitive to specific information (e.g., is the discriminable information for participant *N* only a single factor like memory age, or multiple factors?). The age-selective region was significantly sensitive only to memory age (*t*(29)=4.73, *p*=5.39 × 10^-5^), showing that these voxels only reflect temporal information, and are not sensitive to other factors like memory strength. The people-selective region was sensitive to both people familiarity (*t*(28)=3.02, *p*=0.005) and memory strength (*t*(30)=7.65, *p*=1.57 × 10^-8^). Similarly, the place-selective region was sensitive to both place familiarity (*t*(31)=4.38, *p*=1.25 × 10^-4^) and memory strength (*t*(30)=5.18, *p*=1.40 × 10^-5^). Thus, while these people and place regions are distinct from each other, these regions are also modulated by the strength of the memory. Finally, the memory strength region was sensitive to people familiarity (*t*(28)=2.71, *p*=0.011) and memory strength (*t*(30)=8.16, *p*=4.18 × 10^-9^). We find that the activity in this memory strength mPC region is not predictable by the DNN-based visual features from the videos (see *Methods*, *p*=0.498), suggesting that this region is specifically sensitive to the strength of a memory, and not its visual features.

Finally, we conducted an analysis within the age-specific mPC region to examine the nature of its sensitivity to time. While memory age information has been frequently identified in the hippocampus (Nielson et al., 2015), no such regions have been identified in the mPC to our knowledge. Our whole-brain searchlight RSA suggested that a logarithmic representation of time significantly predicted activity within the mPC, however it is also important to confirm that this region is *specifically* sensitive to a logarithmic representation of time over a linear or exponential representation. Using RSA (see *Methods*), we compared the age-specific mPC subregion to hypothetical models based on logarithmic, linear, or exponential representations of time. We found a small but significant correlation of the mPC subregion with the logarithmic model (Mean *ρ*=-0.006, *t*(31)=2.32, *p*=0.027), but not with the exponential model (Mean *ρ*=0.008, *p*=0.078) or linear model (Mean *ρ*=-0.001, *p*=0.522). The subregion also had a significantly stronger correlation with the logarithmic model than with the exponential model (*t*(31)=2.21, *p*=0.035) or with the linear model (*t*(31)=2.34, *p*=0.026). The negative logarithmic correlation corroborates the results from the whole-brain RSA multiple regression, which observed negative slopes. This suggests that memories that are older or closer in time are represented more distinctively in this brain region.

In sum, these spatially separated peaks across content types suggest a topography of memory information within the mPC, with different clusters representing specific types of information: strength, people, places, and age.

## Discussion

Here, we conducted one of the largest-scale neuroimaging studies of real-world memories, spanning 9,266 total recorded autobiographical videos across 32 visits, from 2 days prior to up to 2,547 days prior (7 years ago). We identified memory-specific signals across the brain that showed greater signal for viewing one’s own memories versus another’s, in spite of identical visual content. We then zoomed in to look at the ability to decode different types of memory properties. We identified effects of memory age and location within the medial temporal lobe, but found that memory strength and emotion could fully account for both of these factors. We also found that memory strength and emotion could be partially determined from stimulus features alone, as measured by an image recognition DNN. However, when we investigated outside the MTL, we identified a region within the mPC with unique variance explained by the age of a memory, with representations of both absolute and relative time. A closer look at the mPC revealed a novel mnemonic topography, with distinct peaks related to memory age, memory strength, people familiarity, and place familiarity. Further, this memory age region within the mPC replicates prior behavioral hypotheses of a logarithmic temporal representation in memory (Singh et al., 2018).

In this study, we were unable to find strong signatures of memory age (or physical distance) in the hippocampus. Recent memories tended to be stronger and more emotional, and both of those factors subsumed any effects of memory age in the hippocampus. This highlights a concern with autobiographical memory research that is often under-addressed: when isolating one property of a memory (e.g., age), it is essential to include all other properties that may be confounded (memory strength, emotion, location, content). Some prior findings of memory recency in the hippocampus could be partially explained by these other confounding factors (e.g., Nielson et al., 2015). However, some other work attempting to factor out the effects of memory strength do still observe higher signal in the hippocampus for recent versus remote autobiographical memories (Gilmore et al., 2021). At first glance, our results appear to support theories that changes in hippocampus signal with memory age instead reflect a weaker memory strength or a semanticization of the memory (Nadel & Moscovitch, 1997; Sekeres et al., 2018). However, higher signal for remote memories (separate from memory strength) in the neocortex supports the predictions of the standard trace model that representations of more remote memories transfer to the neocortex (Alvarez & Squire, 1994). Future studies could investigate specific subfields or subregions of the hippocampus, in case temporal representations exist at finer neural scales. In fact, prior work has identified signals for both recency (Gilmore et al., 2021) and remoteness (Bonnici et al., 2012) specifically in the posterior hippocampus. It is also important to note that these daily recorded memories could elicit differential processing based on type of memory—for example, some videos might elicit recall of a specific episode, while others might elicit a familiarity signal for a routine event. Future research combining memory age with a categorization of these types may reveal a more nuanced story about different theories of consolidation.

However, our study did reveal a key role of the mPC in memory, reflected by a topography of distinct representational clusters for memory age, strength, people, and places. The mPC has only recently emerged as a region for investigating representations of long-term memory content (Brown et al., 2018; Silson et al., 2019a; Woolnough et al., 2020). While the mPC consistently emerges in studies of autobiographical memory, prior accounts hypothesized that the parietal cortex’s involvement in memory was related to working memory or attentional control (Wagner et al., 2005). The parietal cortex was originally believed to play a minor role in autobiographic memory, because early lesion patients were still able to remember events, unlike hippocampal lesion patients (Simons et al., 2008). However, more targeted looks at both medial and parietal lesion patients and healthy volunteers undergoing transcranial magnetic stimulation have revealed that autobiographic memories lose high amounts of detail when the parietal cortex is disrupted (Valenstein et al., 1987; Berryhill et al., 2007; Philippi et al., 2015; Bonnici et al., 2018). The current results reveal a potential role for the mPC in representing detailed content of a retrieved memory across a topography, with separate clusters representing temporal, people, place, and memory strength information. For the memory strength subregion, the precuneus within the mPC has been identified in prior work as a region particularly sensitive to the vividness of recollective memory experiences (Richter et al., 2016; Sreekumar et al., 2018). It is possible that engagement of this subregion could reflect participants performing imagery to assess the vividness of their recollection for those events. Previous work has identified other clusters in the mPC specific to long-term people versus place memory (Silson et al., 2019a), and these clusters are separate from those during perception (Silson et al., 2019b; Bainbridge et al., 2021; Steel et al., 2021). Further, prior work investigating mPC as an “orientation system” in the brain has identified voxel clusters that separately represent comparisons of physical distance, temporal distance, and personal distance, supporting the notion that different regions within the mPC may support different types of information (Peer et al., 2015). Our current study shows that such representations occur beyond the usage of cognitive maps to calculate distance, but also spontaneously during the recall of rich, individual memories from one’s life. However, what specific information is represented in these mPC subregions is still an open question. For example, the subregions that are sensitive to the familiarity of people and places could be some sort of category-specific “familiarity detector” (e.g., Gilmore et al., 2015), could be sensitive to specific types of semantic information associated with these memories, or could contain information about specific familiar items (e.g., face identity). Such questions will be important drivers for future work, and this temporal mPC region we have identified could serve as a key region in which to examine questions about memory consolidation.

While the interactions of memory age, strength, and content were a key aim of our current study, we also investigated the representations of memory location and emotion. Although representations of large spatial scales have been previously identified in the hippocampus (Nielson et al., 2015), we were unable to find clear representations of the spatial location (i.e., geodesic distance) of the memory in medial temporal lobe or medial parietal regions. Given that distant memories tend to be more emotionally positive, contain novel content, and be better remembered, we conjecture that prior findings could have been due to these other characteristics. It is also possible that spatial representations may only occur in the brain at smaller, local scales (e.g., for your neighborhood), and the videos in the current study spanned too broad a set of locations. Indeed, when limiting memories to just 50km around the scan site, some selective regions emerged in the temporal cortex (Supplemental Figure 7), although not in the MTL or mPC. This suggests that future work may need to utilize a constrained set of ranges in order to study geodesic distance. It is also possible that the methods used in the current study were too different from those in prior work (e.g., using recorded videos instead of lifelogging camera snapshots), resulting in divergent findings. We did, however, identify several regions sensitive to the emotion of memory, including the amygdala, medial temporal lobe, visual cortex and many other areas across the cortex. This replicates prior findings supporting an important role for emotion and the amygdala in neural representations of a memory (Piefke et al., 2003; Daselaar et al., 2008), as well as a role of the hippocampus (Ohana et al., 2022). However, unlike prior work that has observed either higher memory strength (Kensinger, 2007) or lower memory strength (D’Argembeau & Van der Linden) for more negative memories, we did not observe a relationship between emotion and memory strength. This lack of relationship could result from how participants used the app; overwhelmingly, participants chose to record more neutral or positive memories over negative memories. There are several possible explanations for this positive memory bias, and a fascinating question is what drives the decision to make a specific memory cue. Perhaps negative memories are often less predictable and also more inappropriate and uncomfortable to record and document. Perhaps participants only want to chronicle positive aspects of their life through this app, especially since these videos are often shared with friends on social media. Or perhaps more generally, the average day tends to be neutral or positive for those who use such apps. There is also the important question of whether such externally recorded videos are truly naturalistic as memory cues, although recent research has shown that mobile videos are effective triggers of memory reactivation even for older adults (Martin et al., 2022). The answer to this question may be changing as our lives become more reliant upon mobile cameras and social media. Future work could examine and even manipulate these different strategies for selecting a mnemonic cue and analyze how they impact representations of these different memories.

The emotion of a memory also showed significant responses in the early visual cortex. This could reflect top-down feedback to V1 (Kravitz et al., 2013; Freese & Amaral, 2005). However, this could also reflect the relationship between the visual features of a memory and its content. In fact, we found that a DNN with similarities to the human visual system (VGG-16; Simonyan & Zisserman, 2014) could predict a participant’s memory strength and emotion for a video based on a single frame. This highlights an important takeaway in memory research— sensory features of a memory may be tightly interlinked with its content and emotion, and some aspects of “memory vividness” could represent visual vividness of the original event. For example, exciting and visually striking events (e.g., a concert, an arcade) could tend to be more memorable and positive. The qualities of a memory itself could also drive the visual features a person chooses to capture in their memory cue. For example, a boring, forgettable day at home could cause a participant to capture a mundane and visually bland video, like a wall or piece of furniture. A memory is thus closely interwoven with the features that represent it, and as memory researchers, we must consider how such sensory features may influence the neural and behavioral representations of such memories. Overall, this study serves as an important demonstration of the complexity of real-world memories—multiple factors such as time, location, strength, emotionality, content, and sensory features are all intertwined, and so these factors must be simultaneously considered even when wanting to only study a single factor.

The current study leveraged a popular mobile app to recruit a set of participants with hundreds to thousands of pre-existing memory stimuli. This resulted in many diverse and representative stimuli over a large time span. While this novel methodology has given us new insights about the neural representations of memories, it also opens up several further questions. Participants had different ranges and sampling of their memories across time, preventing us from looking at the exact same time points across all participants. Future studies could examine more constricted but consistent time points across participants (e.g., having them watch all 365 videos from the last one year). We anticipate such methods could help answer important questions such as the time course at which neocortical consolidation takes place. Participants also differed in their motivations behind using the app, and thus the types of content represented in their videos; for example, some participants recorded similar things gradually changing everyday (e.g., a baby or pet growing up at home), while others intentionally recorded disparate and diverse experiences (e.g., traveling the world or starting college). While we were able to leverage this large variance to simultaneously assess many attributes of memory, some of our effects may be weaker as a result. For example, it is possible that more routine memories would have different representations from distinctive memories, e.g., the former being more gist-like (Koriat et al., 2000; Conway, 2005; Robin & Moscovitch, 2017), and future work could examine how neural representations differ based on the distinctiveness of the memory.

Another key question is the influence of memory reinstatement on representations of these memories. While we limited participants’ reviewing of their memories (see *Methods*), participants could have rewatched some or all memories prior to enrollment in the study. This could influence the level of semanticization of individual memories or the amount of interference across memories. Memory reinstatement is a potential issue with many, if not a majority, of autobiographical memory studies, given that participants often have to supply descriptions of memories or photos to experimenters (triggering reinstatement), and we likely naturally reinstate important memories from our lives. However, future work could quantify the number of replays of memories from an app like 1 Second Everyday to look at influences of reinstatement on memory representations. Interestingly, participants could even be using special strategies to select their memory videos—perhaps intentionally selecting memories that have low interference and rich semantic content. Future studies could have participants utilize such an app with specific constraints, in order to equalize the types of content that are recorded. However, one key advantage of the current method is in our ability to leverage the rich, pre-existing data already collected by the app users. One major obstacle to conducting similar studies with experimental manipulations is that one must build the set of memories from scratch, making it infeasible to look at memories that span multiple years.

The daily images and videos that we record for social media show incredible promise as a new means to examine autobiographical memories. Indeed, future memory work going forward could benefit from a combined examination of both naturalistic observational studies and controlled interventional studies, to benefit from the rich variance and long time scales of the former, but the ability to test targeted hypotheses with the latter. With this rich set of daily recorded memories, we have illustrated the incredible importance of considering the range of factors that make up a memory when measuring their representations in behavior in the brain. Importantly, we have uncovered a topography of mnemonic information in the medial parietal cortex, sensitive to the content, time, and strength of a memory.

## Methods

### The 1 Second Everyday Application

1 Second Everyday is a mobile app available on iOS and Android developed by Cesar Kuriyama in 2013. The app has over 1,500,000 users in the United States, with some users having used the app daily since its inception. The guiding principle of the app is that users should record a brief 1-second video each day, documenting their lives regularly regardless of the excitement or mundaneness of that day. To select their 1-second video for a day, participants record videos of memories throughout the day using the app or their built-in camera app. At the end of the day, the user then selects a 1-second snippet from any of their recorded videos to serve as the second for that day. Thus, a 1-second video represents a user’s choice for the most salient memory cue for that day. An example 1SE video can be viewed on our OSF page at https://osf.io/exb7m/?view_only=8d6939d3fd7c4819aa25aaa5391d10a3. In order to recruit participants for this study, the 1SE app posted an in-app advertisement for users with IP addresses in the Washington, DC area to participate in a study based around their memories.

### Participants

Twenty-three adults (14 female, 9 male) participated in the study, and nine participants returned over 6 months later to participate in a second session. All participants were compensated for their time, and recruitment and experimental procedures were approved by the NIH Institutional Review Board (NCT00001360, 93M-0170). All participants were right-handed, with normal or corrected vision, and had at least 6 months of recorded videos to participate in the study.

Behind the scenes, participants were paired, so that each participant would see videos from their own life as well as videos from another’s life. These participant pairs never met each other; the pairing was only for the purposes of video presentation. When possible, we attempted to pair participants with similar numbers of videos and with similar types of video content (e.g., parents documenting their child’s life; college students). Participants were not given any further information about their paired participant, and were only allowed to see the paired participants’ videos once (in the scanner), to minimize compromising private information of their paired participant. During recruitment, we asked each participant if they knew anyone else interested in the study, to ensure they did not know the person they were paired with. No participant reported recognizing the videos from the other individual.

### Stimuli

App users were recruited for the study if they had at least 6 months of videos recorded. The spread of times included 2 days up to 7 years prior to the day of the scan. Participants supplied us with 762 videos on average (minimum 175, maximum 2210). Participants were asked to provide their videos approximately one week before they were scheduled to come in for the scan. One-second video clips were removed that contained any explicit, obscene, or uncomfortable images (e.g., injuries, spiders, nudity). Clips were also removed that contained any identifiable information of the participants (e.g., full name or address). A black rectangle was placed over the dates embedded in the videos by the app, so participants could not identify the time of the video. In natural use of the app, it is possible that participants watched their own videos multiple times, or different numbers of times per video. In general, the app is usually used to create a compilation of one’s videos once a year (for posting on social media), so we think participants did not repeatedly watch their videos over a long period. Importantly, as soon as participants were recruited for the study (usually 1+ months before coming in for the scan), they were asked not to rewatch any videos they’ve recorded.

For the experiment, we sampled three hundred videos per participant evenly and randomly across their vetted collection of videos. While a majority of participant samples had at least 300 videos (N=23 out of 32), nine participants had fewer than 300 usable videos (170, 250, 260, 260, 270, 280, 286, 294, and 296 videos) and thus all videos were used for their experimental sessions, and we attempted to pair them with participants with smaller numbers of videos. For participants who returned for a second session 6 months later, we collected new videos from them, and sampled a non-overlapping set of videos across the videos provided by them.

While one second is very brief, these videos capture rich amounts of information, and are often highly recognizable to the person who recorded the video. A majority of the time, these videos are outward-facing (not including the recorder) and are selected as a cue to the main activities of that day. Common subjects of a clip may include a concert, an outing with a friend, a favorite TV show, a sports event or game, food, or even boredom at home or one’s office. One second is long enough to identify a location, a song, a sentence, or an event.

Examples of a diverse set of one second videos recorded by the corresponding author can be seen at (https://osf.io/exb7m/?view_only=8d6939d3fd7c4819aa25aaa5391d10a3).

### In-scanner task

Participants first participated in a 7.1-minute localizer task where they saw blocks of faces, objects, scenes, and mosaic-scrambled object images and had to identify a back-to-back repeated image. These images all came from an independent experiment, and the purpose of this scan was to localize face-, scene-, and object-selective visual regions in the participants’ brains. We did not conduct any analyses in the current paper with this localizer data.

Participants then moved onto the main task, which consisted of 10 runs of 6.3 minutes each. In each run, participants saw a set of 60 1-second video clips, 30 of their own and 30 of the partnered participant. Videos were presented with an event-related design, and there was a 5 s inter-stimulus interval between each video. Videos were presented in a randomized order, with their own videos and the partnered participant’s videos randomly intermixed. Importantly, a participant and their paired participant saw the exact same videos in the exact same sequence, so that any differences that emerge are truly due to mnemonic, and not visual, processing of the videos. The videos in each run were selected to evenly sample the entire time span of the participant’s collection of videos. For example, if a participant had 10 months of videos, Run 1 would contain 30 videos spanning the full 10 months (3 videos per month), as would Runs 2, 3, etc. This even sampling was performed with a random date jitter, to avoid embedding structure in the sampling (e.g., avoiding always sampling from the same day of the week). Participants with fewer videos experienced 10 scanner runs that each showed fewer videos and spanned a shorter period of time.

While watching the videos, participants performed a visual imagery task. They were instructed that when they saw one of their own videos, they should try to remember what was occurring in that video and surrounding memories for that day. When they saw someone else’s videos, they were instructed to imagine what was occurring in that video and its surrounding context. Participants did not make any button presses during the task. All in-scanner tasks were coded in Psychtoolbox-3 for MATLAB (Brainard, 1997).

### Post-scan video labeling

After the scan, participants completed a behavioral task in which they saw each of their own ∼300 videos and labeled them for a wide range of information. Participants first identified where the video occurred, as specifically as possible, using a Google Maps API-based search that saved the latitude and longitude of the location. This was generally relatively easy for participants, as they could input addresses, business names (e.g., a restaurant name), or a business name combined with location (e.g., the Starbucks in Bethesda). Participants rated memory strength of the videos (“how strong is your memory for this event?”) on a Likert scale of 1 (very weak) to 5 (very strong). They then indicated the length of the memory that 1-second cue retrieved: seconds, minutes, hours, the whole day, or N/A (if they did not retrieve a memory); this data was not analyzed for the current study. They labeled the emotions for the event (“how would you describe your emotions for this event?”): Very negative, somewhat negative, neutral, somewhat positive, and very positive. They also measured the content of the memory: 1) whether it contained people and how familiar they were with them at the time of the memory, 2) whether it contained the participant themself, and 3) whether the place depicted in the video was new or familiar at the time of the memory. Finally, to quantify any flashbulb memories, we asked them to name videos that contained an important event (not analyzed in this study). Participants could work on this labeling task remotely, as it usually took 2-4 hours to complete, but were asked to complete it within 48 hours of the scan. They were not allowed to search for information or ask others for assistance in remembering an event.

### Scanning parameters

The experiment was conducted at the NIH Clinical Center, using a 3 Tesla General Electric MRI scanner system with a 32-channel head coil. Whole-brain anatomical scans were acquired using the MP2RAGE sequence, with 1mm isotropic voxels. Whole-brain functional scans were acquired with an EPI scan of 2.5mm isotropic voxels (repetition time = 2500ms, echo time = 30ms, flip angle = 75 degrees), with slices aligned parallel to the temporal lobe. Functional scans were pre-processed with slice timing correction and motion correction using AFNI and surface-based analyses were performed using MATLAB and SUMA (Cox, 1996; Saad and Reynolds, 2012). No smoothing was applied. Anatomical regions of interest (ROI) were localized in the MTL including the hippocampus, PHC, ERC, and amygdala, using FreeSurfer’s *recon-all* pipeline. Group contrasts were generated in the surface-space by surface-based alignment to a template surface, and in the volume space by alignment to the MNI template space.

### Analyses

#### General linear model

First, in order to observe differences between viewing one’s own videos versus another’s videos, we conducted a general linear model (GLM) in each participant, with videos split between two conditions: their own videos, and another’s videos. Group univariate contrasts were then produced using a T-contrast of own vs. other’s videos. We conducted the group contrast across all participant samples (Figure 4, Supplemental Figure 2), as well as just within the first and second sessions of the nine participants who came in for two sessions separated by at least 6 months (Supplemental Figure 4).

Next, to model the voxel values for each video, we conducted a GLM that included every video as a 6-second block regressor convolved with a canonical hemodynamic response function. The t-statistic of the beta value for a given video was taken as its voxel value for all following analyses. For both GLMs, six nuisance regressors were also included for motion (x, y, z, roll, pitch, yaw), computed from the motion correction. The resulting beta values for each video were then utilized as inputs for the proceeding fMRI analyses: the regression analyses, the representational similarity analyses, and the voxel peaks comparison.

#### Regression analyses for relating mnemonic content and the brain

We employed regressions to assess the degree to which various aspects of a memory were related to the voxel values within a given region. We conducted these analyses first within our anatomical ROIs in the MTL. We then conducted these same analyses in a whole-brain searchlight of 5-voxel diameter iteratively moved throughout the brain. The main predictors modeled for every video and entered into the regression were:

(1) *Memory age*, calculated as the number of days the memory occurred before the scan.
(2) *Memory strength*, the participant’s rating of their strength of the memory (1 = very weak, to 5 = very strong).
(3) *Emotion*, the participant’s rating of their emotions for the memory (1 = very negative, to 5 = very positive)
(4) *Memory distance*, calculated as the geodesic distance of the memory’s location to the scanning center at the NIH.

These predictors were then entered into a model to predict the rank-ordered mean voxel value (t-statistic) within the ROI for each video. These t-statistics were rank-transformed in order to avoid any assumptions of a linear relationship (given that time may vary logarithmically). For example, we can test whether older memories corresponded with higher (or lower) voxel activation without assuming the voxel activation itself should necessarily increase linearly (just the ranking). To observe the relationships between a single predictor and the ranked voxel value, these were computed as simple linear regressions (where *β* is the resulting slope, and *ε* is the residual and intercept):

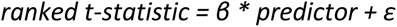

We then computed multiple regressions to examine the contribution of each predictor to the mean voxel value. No predictors showed evidence of multicollinearity (all *r* < 0.20, in contrast with a rule of thumb of *r* > 0.80 indicative of multicollinearity).

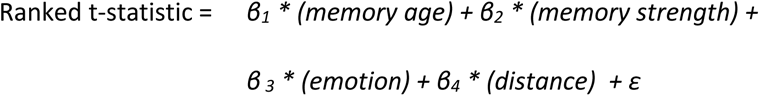

The resulting betas across participants were then compared against a null hypothesis slope of 0 using a one-sample two-tailed t-test. If the average slope (beta) across participants for a given predictor is significantly different from 0, this suggests a significant positive (or negative) relationship between the predictor and voxels in the brain. We report Cohen’s *d* for effect size, and performed FDR correction (*q*<0.05) across the ROIs for each analysis. We ran this multiple regression both for predicting the voxel values for watching one’s own videos from one’s own ratings of those videos, but also for predicting the voxel values for watching another’s videos from the other’s ratings of those videos.

#### Representational similarity analyses of memory content

To serve as a complementary measure to the univariate regression analysis, we also analyzed patterns of memory content in the brain using representational similarity analysis (RSA). RSA tests the degree to which behavioral hypotheses relate with patterns in the brain, by observing the similarity between pairs of stimuli (Kriegeskorte et al., 2008). The idea is that, within a given brain region, the similarity of patterns of voxels across stimuli can capture the representational geometry of that region; for example, stimuli with more similar patterns may also be more cognitively similar. The voxel-based representational geometry of a region can thus be compared to hypothetical representational geometries of behavioral measures to test the involvement of different cognitive processes. Representational geometries are compared as representational dissimilarity matrices (RDMs), or matrices representing the pairwise distances between all stimuli. Here, neural distance in a given region between two memories was calculated as 1 minus the Pearson correlation of the voxel vectors of each memory. Behavioral distances (e.g., distance in memory age) between two memories were calculated as the absolute value of the difference in the behavioral measure for each memory. RDMs were thus calculated from the brain and the behavioral measures of memory strength, emotion, memory age, and distance.

We conducted RSA in the anatomical ROIs in the MTL, as well as across the whole brain. For both, we tested a multiple regression to observe the degree to which the behavioral RDMs in conjunction could predict the brain-based RDM. Time and distance were modeled logarithmically, based on prior work suggesting logarithmic representations of time and distance using a similar multiple regression RSA framework in the hippocampus (Nielson et al., 2015). For memory age, we took the base-10 log of each measure of memory age (number of days from scan) before creating the memory age RDM. For memory distance, we modeled each memory as a vector with the start point at the scan site and the endpoint at the memory location (defined by latitude and longitude indicated by the participant). For each pair of memories, we took the base-10 log of the magnitude of each pair of vectors. We then calculated the Euclidean distance between their endpoints. These log-transformed pairwise distances were then used as the distance measures in the memory distance RDM. Memory strength and emotion were kept in their original units (not log-transformed), given that they were rated on a Likert scale with no specific prediction of a logarithmic representation. The final regression formula tested in each region was as follows:

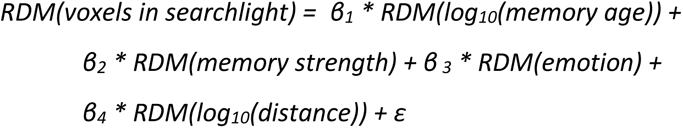

Group maps were then formed by taking the average beta across participants for a given predictor. Significance was tested using t-tests comparing the beta across participants to a null hypothesis of 0. Whole-brain analyses were conducted in searchlights across the whole brain with a diameter of 5 voxels.

#### Representational similarities of temporal representations

While our whole-brain searchlight RSA suggests a logarithmic representation of time in the memory age mPC region, we wanted to confirm that time was indeed best modeled logarithmically. To this end, we conducted RSA within the memory age mPC ROI to compare different hypothetical representations of memory age. This ROI was defined as the significant mPC region identified by the multiple regression group analysis (Figure 6), transformed into each participant’s native space. We tested three RSA models based on hypotheses about memory age. These models were constructed as matrices, where each cell contains the pairwise temporal distance between two memories. First, we tested a linear hypothesis where the distance between two memories is constant regardless of how remote they are. Pairwise distance was calculated as *|timeB – timeA|*. Second, we tested a logarithmic hypothesis where the temporal distance between two memories decreases as they become more remote.

Pairwise distance was calculated as *|log10(timeB) – log10(timeA)|*. Third, we tested an exponential hypothesis where the temporal distance between two memories exponentially increases as they become more remote. Pairwise distance was calculated as *|10^timeB^ – 10^timeA^|*.

In order to compare each hypothesis with the brain, a RDM was constructed for the ROI. For each video, we extracted its vector of voxel activations in that ROI. We then constructed a matrix of the 1 – the Pearson correlation between all pairs of memory voxel vectors. To compare the brain to these temporal hypotheses, we then took the Spearman rank correlation between the upper triangle of this ROI-based matrix with the upper triangle of each temporal hypothesis matrix. This Spearman correlation was calculated for every participant, and then Fisher Z-transformed. Finally, these transformed correlation coefficients were compared to a null hypothesis of 0 with a one-sample t-test, and compared to each other using paired-samples t-tests, to see whether any one temporal hypothesis better explained the data than the others.

#### Computer vision predictions of video memory strength and emotion

We conducted two analyses to test the relationship of the visual features within a video and ratings of memory strength and emotion: 1) an analysis based on a visual DNN, and 2) an analysis based on optical flow. To extract the visual features for each memory, we took the middle frame of each 1-second video, and inputted it into the VGG-16 DNN for object classification (Simonyan & Zisserman, 2014). The early layers of VGG-16 are typically thought to extract early visual features of an image (e.g., color, edges, orientations), while the later layers of VGG-16 are thought to extract late visual and semantic information (e.g., specific object parts and categories, like faces). These visual DNNs have also shown correspondences with the human visual system (Yamins et al., 2014; Cichy et al., 2016). The VGG-16 DNN outputted a vector of values at each layer for each middle frame of the videos. We then tested the degree to which the early and late layer output vectors can predict a person’s rating of memory strength and emotion for a video. For the early layer, we looked at layer 2 (the first layer after the image input), while for the late layer, we looked at layer 20 (the penultimate layer before the category prediction). For a given participant, 80% of their videos were utilized as a training set for a support vector regression (SVR) that learned to predict memory strength ratings from a given layer’s outputs for a given video’s middle frame. We then tested this model on the remaining held-out 20% of their videos, and correlated memory strength predictions with actual memory strength. This training and testing was repeated iteratively across 25 iterations per participant, resulting in an average correlation coefficient reflecting prediction performance. The same analysis pipeline was also conducted for predicting a video’s emotion rating from the early and late layer outputs of the video’s middle frame.

We also tested whether VGG-16 could predict brain activity in the memory strength subregion in the mPC. We ran a similar analysis, where for each participant we conducted 25 iterations of a SVR, using 80% of the data for training and a hold-out of 20% of the data for testing the model. For each iteration, an SVR was trained to predict the mean brain activation in the group-defined memory strength mPC subregion, based on the layer 20 values from VGG-16. Prediction performance was calculated as the average Pearson correlation between the predicted and actual brain activation measures for the held-out test data across the 25 iterations.

Next, we examined the relationship of motion to memory strength and emotion. We estimated motion using a toolbox for real-time optical flow (Karlsson & Bigun, 2019), which conducts edge detection across an image and then estimates movement of those edges using the Lucas-Kanade method (Lucas & Kanade, 1981). For a given one-second clip, we calculated the total magnitude of optical flow between the first and middle frame, and between the middle and last frame of the video, and then averaged these two measures. This provides a measure of how much motion there is in the video. For each participant, we then calculated a Spearman’s rank correlation between the amount of motion in a video and its memory strength ratings or emotion ratings to assess how much motion was predictive of these features.

To test the success of these predictions across participants, we compared the mean correlation coefficients across participants against a null hypothesis of 0. Because these correlation coefficients may not be normally distributed, they were compared utilizing Wilcoxon sign rank tests, which test against a median of 0 (rather than a mean of 0). A significant positive result indicates that prediction accuracy is significantly higher than chance.

#### Voxel peaks comparison across memory factors

In order to compare the distributions of memory information in the mPC, we visualized the peaks of significant activation for different types of information. First, the significant voxels for each information type were identified using group-level univariate contrasts. People familiarity voxels were identified with a T-contrast of memories with familiar people versus novel people. Three participants did not have at least five videos with novel people, and so were excluded from this analysis. Place familiarity voxels were identified with a T-contrast of memories occurring within familiar versus novel places. Memory age voxels were identified with a T-contrast of recent memories (within the last three months) and remote memories (more than three months ago). This cut-off was selected so that there would be a large number of memories in each bin, while trying to avoid the “recent” bin containing too many older memories (thus potentially washing out any effects). However, similar peaks emerge for other time cut-offs. Memory strength voxels were identified using an ANOVA distinguishing the five memory strength response options. One participant was excluded from this analysis for not having at least five weak memories. Finally, we also investigated the top 1000 voxels for an ANOVA distinguishing the five emotion response options, however no voxels occurred in the mPC (Supplemental Figure 11). All contrasts were thresholded at *p*<0.05, and then we visualized the top 1000 of these voxels to identify peak regions associated with each property of memory.

To test the selectivity of these regions, we conducted a leave-one-out analysis at the participant level. Participants were aligned to the surface space. For each participant, we identified the top 1000 voxels for each group contrast (memory age, people familiarity, place familiarity, memory strength) as described above, with all other participants. We then took the medial parietal portion of each contrast, defined as any of these significant voxels on the medial surface posterior to the central sulcus. Within each of these mPC regions defined by *N*-1 participants, for the excluded participant *N* we then measured the mean beta value of the four contrasts (memory age, people familiarity, place familiarity, and memory strength). Essentially, this allows us to see whether a set of voxels sensitive to a certain type of information (e.g., memory age) in all other participants shows this same sensitivity in the left-out participant. This was repeated for all 32 participant samples. Note that no constraints were put on the regions, so they could include overlapping voxels. To test the selectivity of each region, we then conducted two-tailed t-tests of the measures for the 32 samples versus a null hypothesis of 0. The resulting statistics were then FDR corrected with a threshold of *q*<0.05.

### Data Availability

Video features, memory features, and fMRI data will be publicly available at an Open Science Framework repository: https://osf.io/exb7m/?view_only=8d6939d3fd7c4819aa25aaa5391d10a3. Raw memory videos and memory geocoordinate information will not be made available, to protect the personally identifiable information of the participants.

## Acknowledgements

Big thanks to Cesar Kuriyama for allowing us to recruit participants through his app for this study. Thank you to Matthias Nau and Adrian Gilmore for their invaluable advice on the manuscript. Thank you also to Wan Kwok, Elizabeth Hall, Anna Corriveau, Alexis Kidder, and Adam Dickter for their help in data collection and participant recruitment on this study. Thank you to Seyed-Mahdi Khaligh-Razavi, Aude Oliva, Jesse Rissman, and Marc Howard for their advice on early versions of this project. This research was supported by the Intramural Research Program of the National Institutes of Health (ZIA-MH-002909), under National Institute of Mental Health Clinical Study Protocol 93-M-1070 (NCT00001360).

