## Supplemental Information for "Multidimensional topography of memory revealed from thousands of daily documented memories"

Beecher Hall 303

Chicago, IL 60637

**Table of Contents**

pages

3	**Supplemental Figure 1.** Distribution of participant’s videos, as categorized by their locations and people

4	**Supplemental Figure 2.** Unthresholded activation differences based on video mnemonic content

5	**Supplemental Figure 3.** People and place familiarity ROIs

6	**Supplemental Figure 4.** Mnemonic contrasts for the two separate experimental sessions

7	**Supplemental Table 1.** Comparisons of memory properties and ROI signal by hemisphere

8	**Supplemental Table 2.** Comparisons of memory properties in a combined model and ROI signal

9	**Supplemental Table 3.** Comparisons of memory properties and ROI signal for another’s videos

10	**Supplemental Figure 5.** Alternate views of memory content representations

11	**Supplemental Figure 6.** Representations of different memory content, at stringent thresholds

12	**Supplemental Figure 7.** Map of memory content representations, for a constrained set of distances (within 50 km)

13	**Supplemental Figure 8.** Map of memory content representations, with regressors included for people and place familiarity

14	**Supplemental Figure 9.** Whole-brain views of the memory topography

15	**Supplemental Figure 10.** Whole-brain views of the memory topography on the same colormap

16	**Supplemental Figure 11.** Whole-brain views of the top voxels for emotion

17	**Supplemental Figure 12.** Top voxels for people and place familiarity

a)
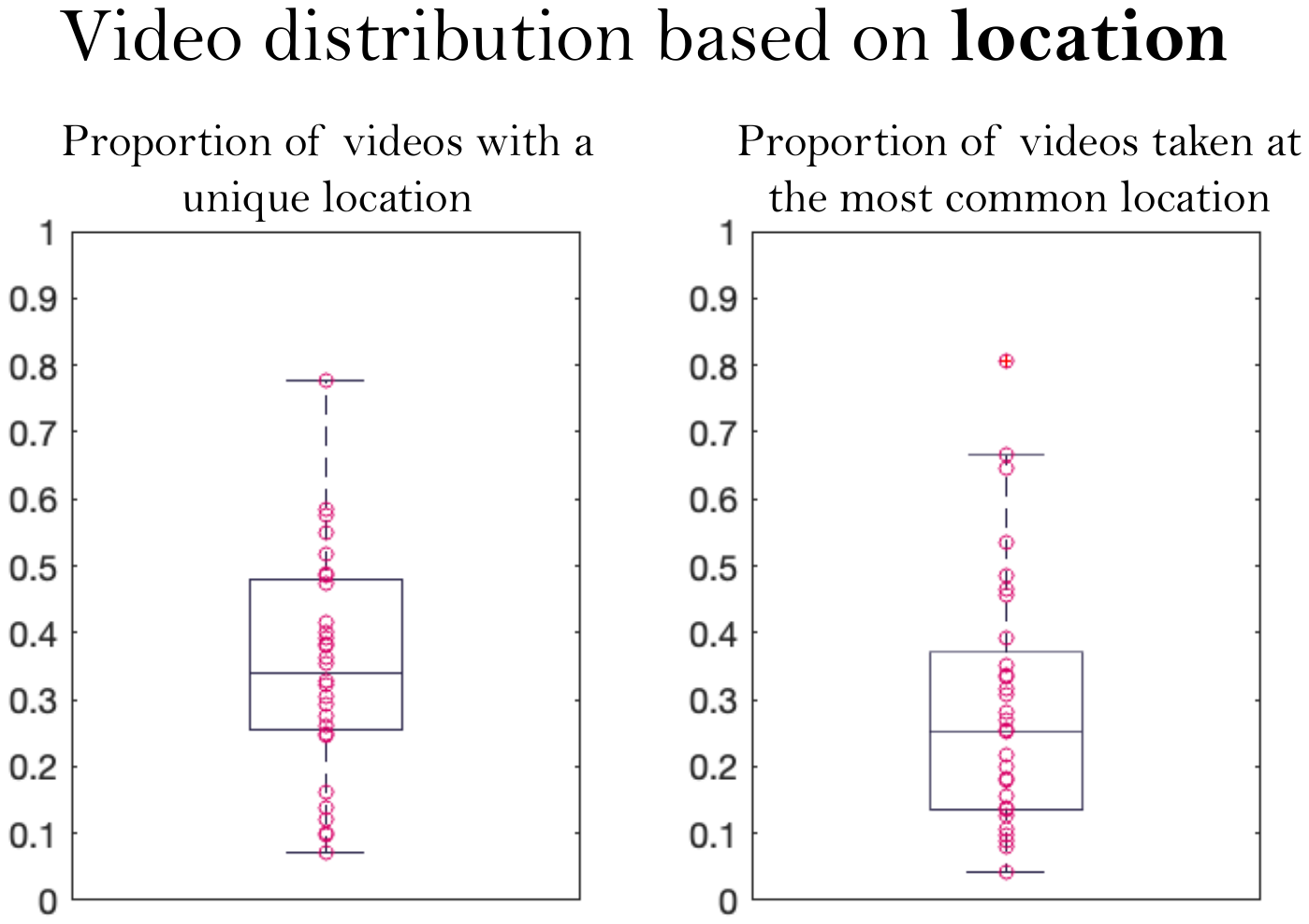

b)
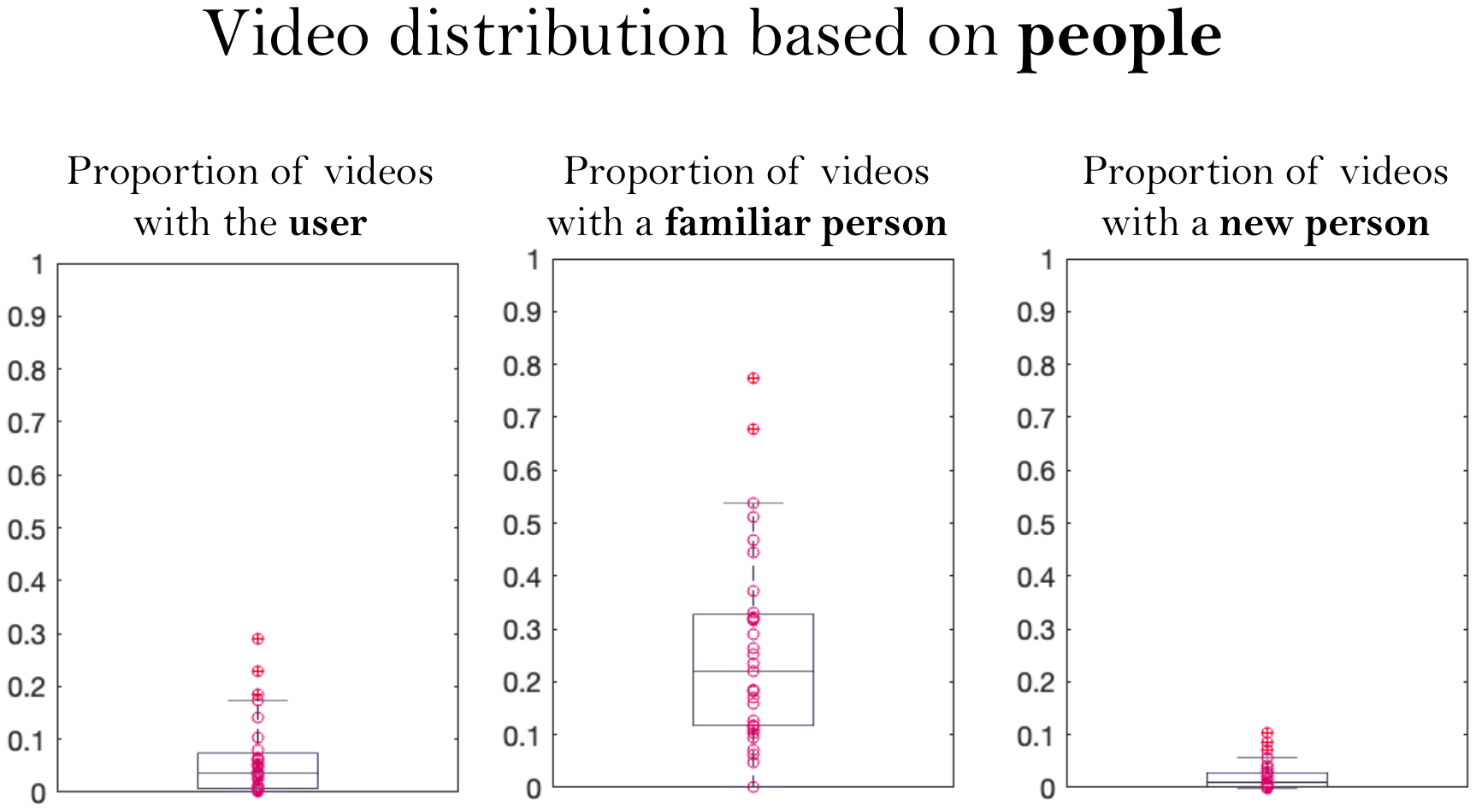

**Supplemental Figure 1. Distributions of participant’s videos, as categorized by their locations and people.** (a) Scatterplots and boxplots where each dot represents the proportion of videos occurring in a unique location (left) and the most common location (right) for all participants. (b) Scatterplots and boxplots where each dot represents the proportion of videos containing the user in the video (a “selfie”; left), the proportion of videos containing a familiar person in the video (middle), and the proportion of videos containing a novel person in the video (right).

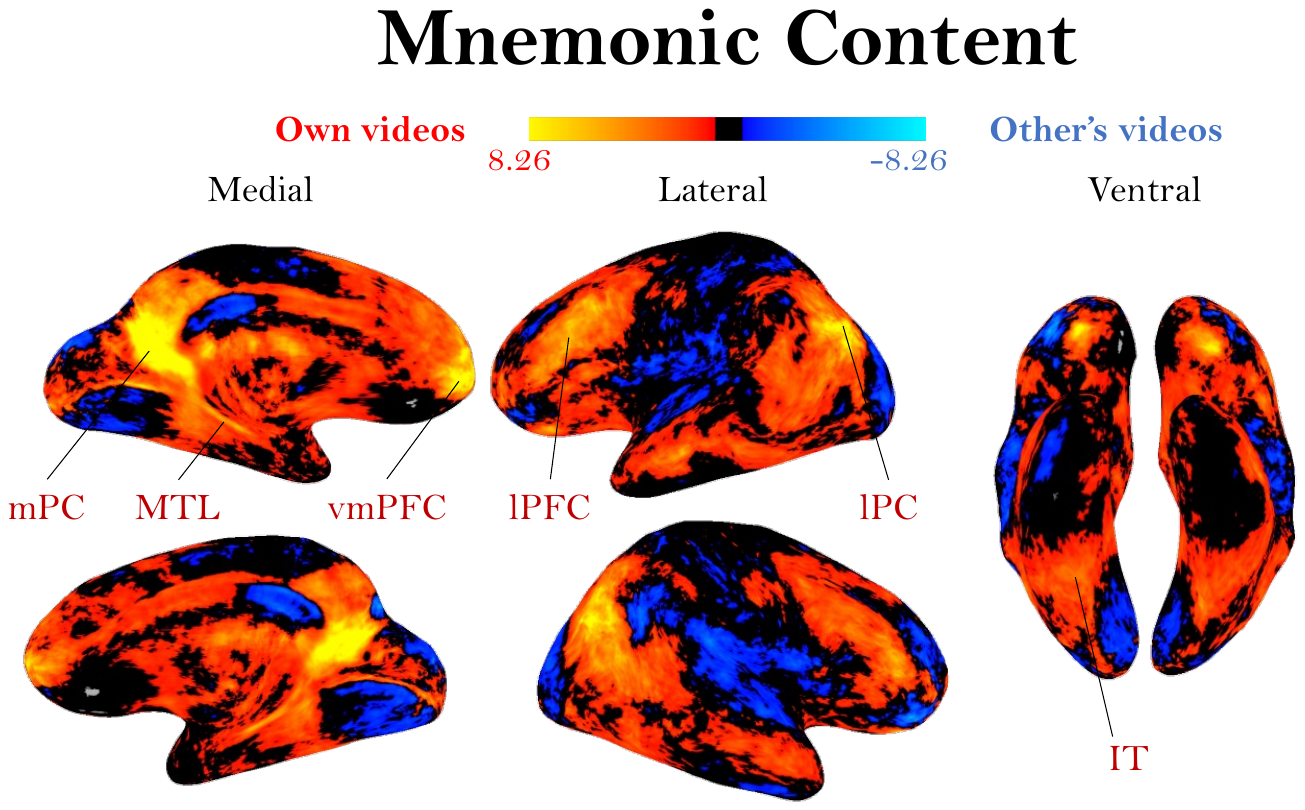

**Supplemental Figure 2. Unthresholded activation differences based on video mnemonic content.** This figure is an unthresholded version of Figure 4 in the manuscript, showing a whole-brain group activation map (N=32) for participants viewing their own videos (red/yellow) versus viewing the other person’s videos (blue). The colormap represents the range of beta values.

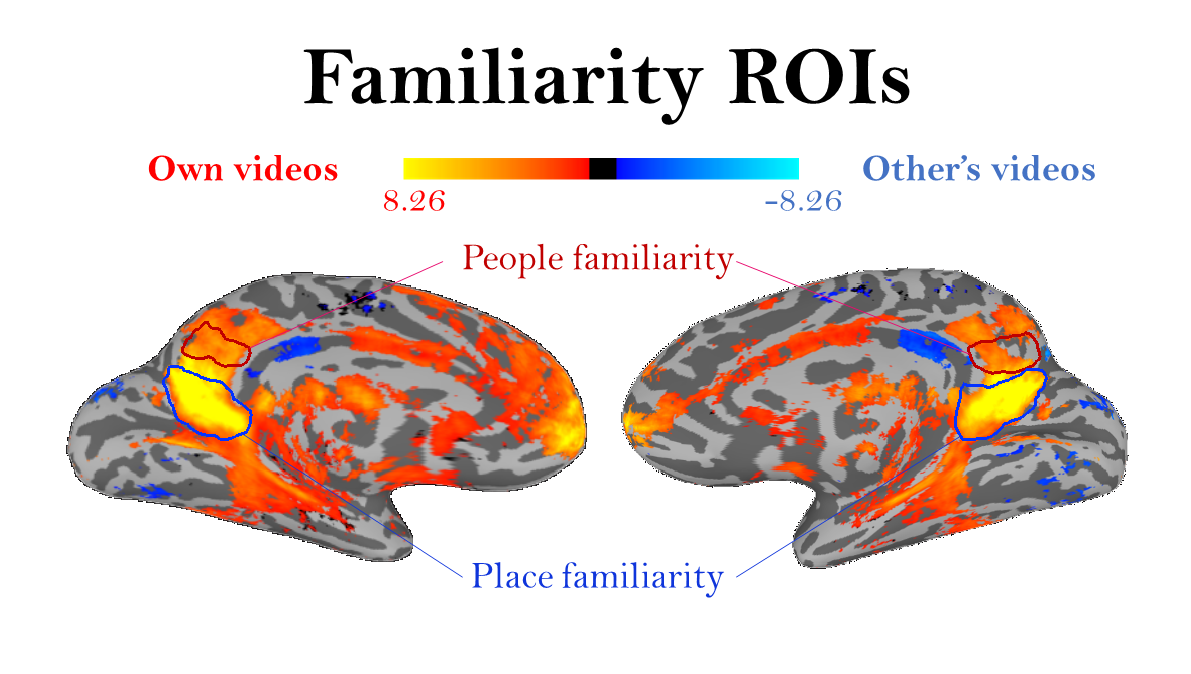

**Supplemental Figure 3.** **People and place familiarity ROIs.** Depictions of the main medial parietal ROIs identified in Silson et al. (2019a) that are selective for familiar people over familiar places (in red), and for the opposite contrast of familiar places over familiar people (in blue). The colormap represents the range of beta values. These ROIs are overlaid on the contrast shown in Figure 4 in the main manuscript (N=32, whole-brain group activation for viewing one’s own videos versus the other participant’s videos).

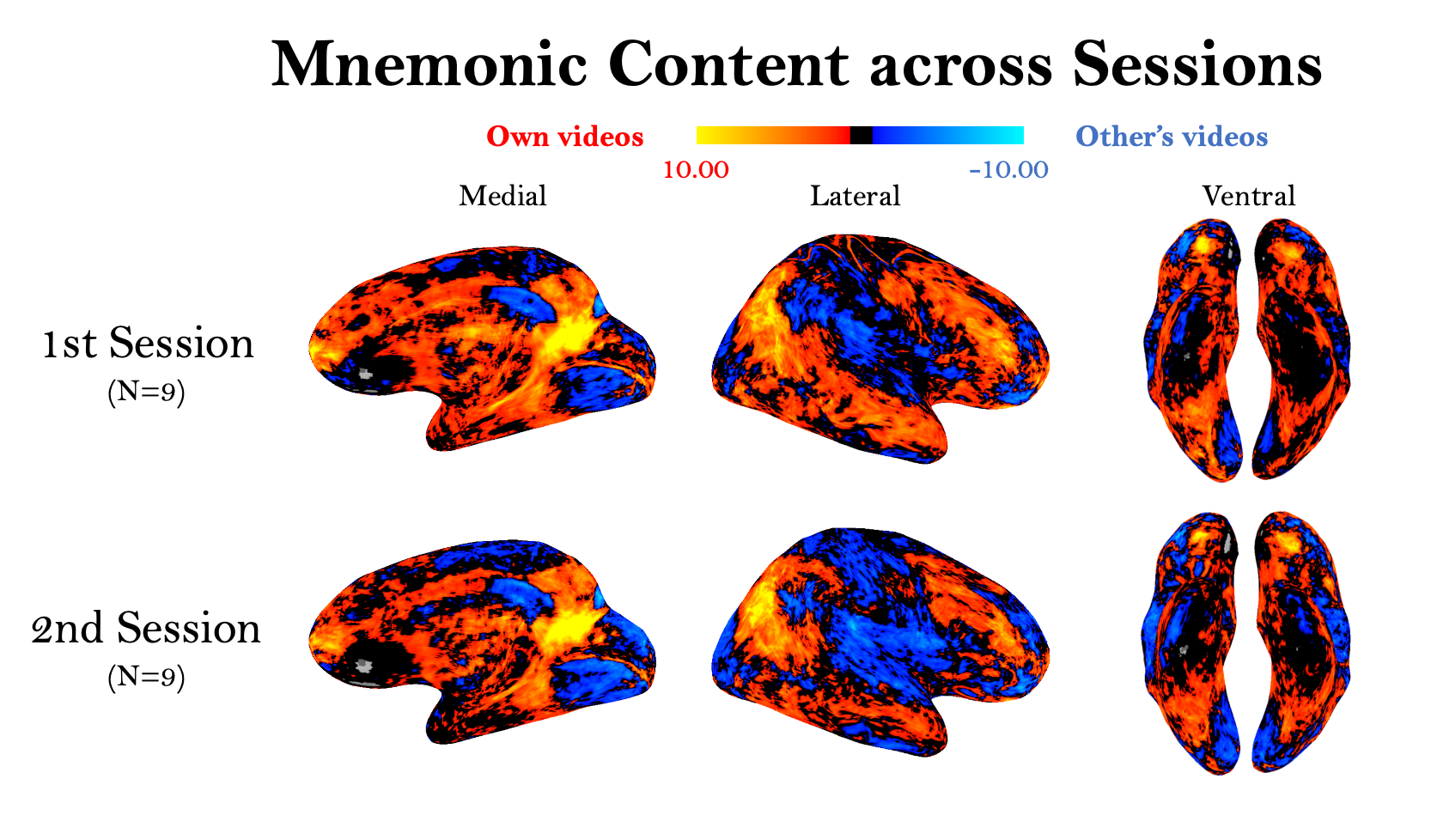

**Supplemental Figure 4.** **Mnemonic contrasts for the two separate experimental sessions.** Nine individuals participated in the experiment in two sessions, at least six months apart and containing no overlapping videos. These maps show the contrast of viewing one’s own videos versus another participants’ videos (as in Figure 4 and Supplemental Figure 2) for those participants using data from their first session only (top) and their second session only (bottom). The colormap represents the range of beta values. Despite encompassing different sets of videos, the regions that emerge based on memory for one’s own videos remain largely the same.

**Supplemental Table 1. Comparisons of memory properties and ROI signal by hemisphere.** Shown are results from simple linear regressions quantifying the relationship of a given memory property (age, memory strength, emotion, and distance) and mean beta value, across voxels in anatomically-defined ROIs within the medial temporal lobe. Indicated are the beta-value (*β*) from the regression, and the p-value resulting from the t-test comparing the beta values across participants with a null hypothesis of 0 slope. Regressions with *p* < 0.05 are colored in grey. Age is measured as number of days from the scan; memory strength is from 1 (very weak) to 5 (very strong); emotion is from 1 (very negative) to 5 (very positive); and distance is number of km from the scanning center. (Hipp = hippocampus, Amyg = amygdala, ERC = entorhinal cortex, PHC = parahippocampal cortex.)

|  | **Memory age** | | **Memory strength** | | **Emotion** | | **Distance** | |
| --- | --- | --- | --- | --- | --- | --- | --- | --- |
| ROI | *β* | *p* | *β* | *p* | *β* | *p* | *β* | *p* |
| L-Hipp | -0.021 | 0.006 | 6.184 | 1.42E-5 | 6.545 | 1.89E-4 | 0.004 | 0.255 |
| R-Hipp | -0.018 | 0.028 | 6.181 | 4.86E-6 | 6.641 | 4.70E-5 | 0.003 | 0.18 |
| Hipp | -0.019 | 0.009 | 6.183 | 4.46E-6 | 6.593 | 7.27E-5 | 0.003 | 0.209 |
| L-Amyg | -0.019 | 0.024 | 5.719 | 8.11E-5 | 7.085 | 4.23E-5 | 0.004 | 0.276 |
| R-Amyg | -0.022 | 0.018 | 5.712 | 4.46E-6 | 7.236 | 3.59E-5 | 0.004 | 0.011 |
| Amyg | -0.021 | 0.013 | 5.716 | 1.09E-5 | 7.161 | 2.18E-5 | 0.004 | 0.073 |
| L-ERC | -0.022 | 0.028 | 5.311 | 4.49E-6 | 6.374 | 2.81E-5 | -3.74E-4 | 0.88 |
| R-ERC | -0.023 | 0.005 | 5.074 | 2.10E-5 | 5.943 | 1.35E-4 | -2.11E-4 | 0.939 |
| ERC | -0.023 | 0.010 | 5.192 | 6.37E-6 | 6.158 | 4.52E-5 | -2.93E-4 | 0.909 |
| L-PHC | -0.023 | 0.008 | 4.716 | 7.40E-5 | 4.096 | 0.006 | 0.001 | 0.668 |
| R-PHC | -0.022 | 0.004 | 4.168 | 1.82E-4 | 3.615 | 0.018 | 0.004 | 0.024 |
| PHC | -0.023 | 0.005 | 4.442 | 8.10E-5 | 3.856 | 0.008 | 0.003 | 0.212 |

**Supplemental Table 2. Comparisons of memory properties in a combined model and ROI signal.** Shown are results from a multiple linear regression quantifying the relationship of memory properties in a combined model and mean beta value, across voxels in anatomically-defined ROIs within the medial temporal lobe. Indicated are the beta-value (*β*) from the regression, and the p-value resulting from the t-test comparing the beta values across participants with a null hypothesis of 0 slope. Regressors with *p* < 0.05 are colored in grey. Age is measured as number of days from the scan; memory strength is from 1 (very weak) to 5 (very strong); emotion is from 1 (very negative) to 5 (very positive); and distance is number of km from the scanning center. (Hipp = hippocampus, Amyg = amygdala, ERC = entorhinal cortex, PHC = parahippocampal cortex.)

|  | **Memory age** | | **Memory strength** | | **Emotion** | | **Distance** | |
| --- | --- | --- | --- | --- | --- | --- | --- | --- |
| ROI | *β* | *p* | *β* | *p* | *β* | *p* | *β* | *p* |
| L-Hipp | -0.011 | 0.22 | 4.681 | 1.71E-4 | 3.728 | 0.008 | -0.002 | 0.669 |
| R-Hipp | -0.004 | 0.712 | 4.709 | 5.87E-4 | 3.931 | 0.012 | -0.003 | 0.387 |
| Hipp | -0.008 | 0.411 | 4.695 | 1.57E-4 | 3.829 | 0.009 | -0.003 | 0.534 |
| L-Amyg | -0.007 | 0.442 | 3.925 | 0.001 | 5.064 | 1.42E-4 | -0.001 | 0.755 |
| R-Amyg | -0.006 | 0.578 | 3.915 | 0.001 | 4.866 | 0.002 | -0.002 | 0.491 |
| Amyg | -0.006 | 0.474 | 3.920 | 6.12E-4 | 4.965 | 4.59E-4 | -0.002 | 0.637 |
| L-ERC | -0.013 | 0.211 | 3.578 | 0.002 | 4.541 | 0.002 | -0.005 | 0.178 |
| R-ERC | -0.011 | 0.220 | 3.294 | 0.008 | 3.944 | 0.007 | -0.006 | 0.242 |
| ERC | -0.012 | 0.194 | 3.436 | 0.003 | 4.242 | 0.003 | -0.005 | 0.209 |
| L-PHC | -0.012 | 0.182 | 3.345 | 0.007 | 1.772 | 0.185 | -0.003 | 0.430 |
| R-PHC | -0.014 | 0.123 | 2.728 | 0.035 | 1.724 | 0.289 | -5.89E-4 | 0.776 |
| PHC | -0.013 | 0.134 | 3.037 | 0.014 | 1.748 | 0.218 | -0.002 | 0.530 |

**Supplemental Table 3. Comparisons of memory properties and ROI signal for another’s videos.** Shown are results from a multiple linear regression quantifying the relationship of memory properties in a combined model and mean beta value for viewing videos recorded by another participant, for which the current participant has no memory. Regressions were run across voxels in anatomically-defined ROIs within the medial temporal lobe. Indicated are the beta-value (*β*) from the regression, and the p-value resulting from the t-test comparing the beta values across participants with a null hypothesis of 0 slope. Regressions with *p* < 0.05 are colored in grey. Age is measured as number of days from the scan; memory strength is from 1 (very weak) to 5 (very strong); emotion is from 1 (very negative) to 5 (very positive); and distance is number of km from the scanning center. These measures all came from the participant who recorded the videos, not the one viewing them in the scanner. (Hipp = hippocampus, Amyg = amygdala, ERC = entorhinal cortex, PHC = parahippocampal cortex.)

|  | **Memory age** | | **Memory strength** | | **Emotion** | | **Distance** | |
| --- | --- | --- | --- | --- | --- | --- | --- | --- |
| ROI | *β* | *p* | *β* | *p* | *β* | *p* | *β* | *p* |
| L-Hipp | 0.005 | 0.518 | 2.042 | 0.025 | 2.039 | 0.187 | -0.002 | 0.274 |
| R-Hipp | 0.01 | 0.300 | 1.848 | 0.060 | 2.246 | 0.156 | -0.007 | 0.121 |
| Hipp | 0.008 | 0.321 | 1.945 | 0.030 | 2.142 | 0.152 | -0.005 | 0.092 |
| L-Amyg | 0.008 | 0.289 | 1.414 | 0.124 | 2.751 | 0.092 | -0.003 | 0.320 |
| R-Amyg | 0.006 | 0.522 | 2.248 | 0.047 | 2.059 | 0.184 | -0.009 | 0.099 |
| Amyg | 0.007 | 0.361 | 1.831 | 0.050 | 2.405 | 0.101 | -0.006 | 0.078 |
| L-ERC | 0.006 | 0.353 | 1.755 | 0.052 | 1.181 | 0.444 | -0.010 | 0.138 |
| R-ERC | 0.004 | 0.553 | 1.325 | 0.153 | 0.39 | 0.806 | -0.003 | 0.471 |
| ERC | 0.005 | 0.405 | 1.54 | 0.080 | 0.785 | 0.597 | -0.007 | 0.138 |
| L-PHC | 0.006 | 0.404 | 2.279 | 0.040 | -1.282 | 0.376 | 2.65E-4 | 0.968 |
| R-PHC | 0.007 | 0.534 | 1.774 | 0.076 | -0.698 | 0.615 | 0.001 | 0.812 |
| PHC | 0.006 | 0.445 | 2.027 | 0.047 | -0.99 | 0.468 | 7.05E-4 | 0.900 |

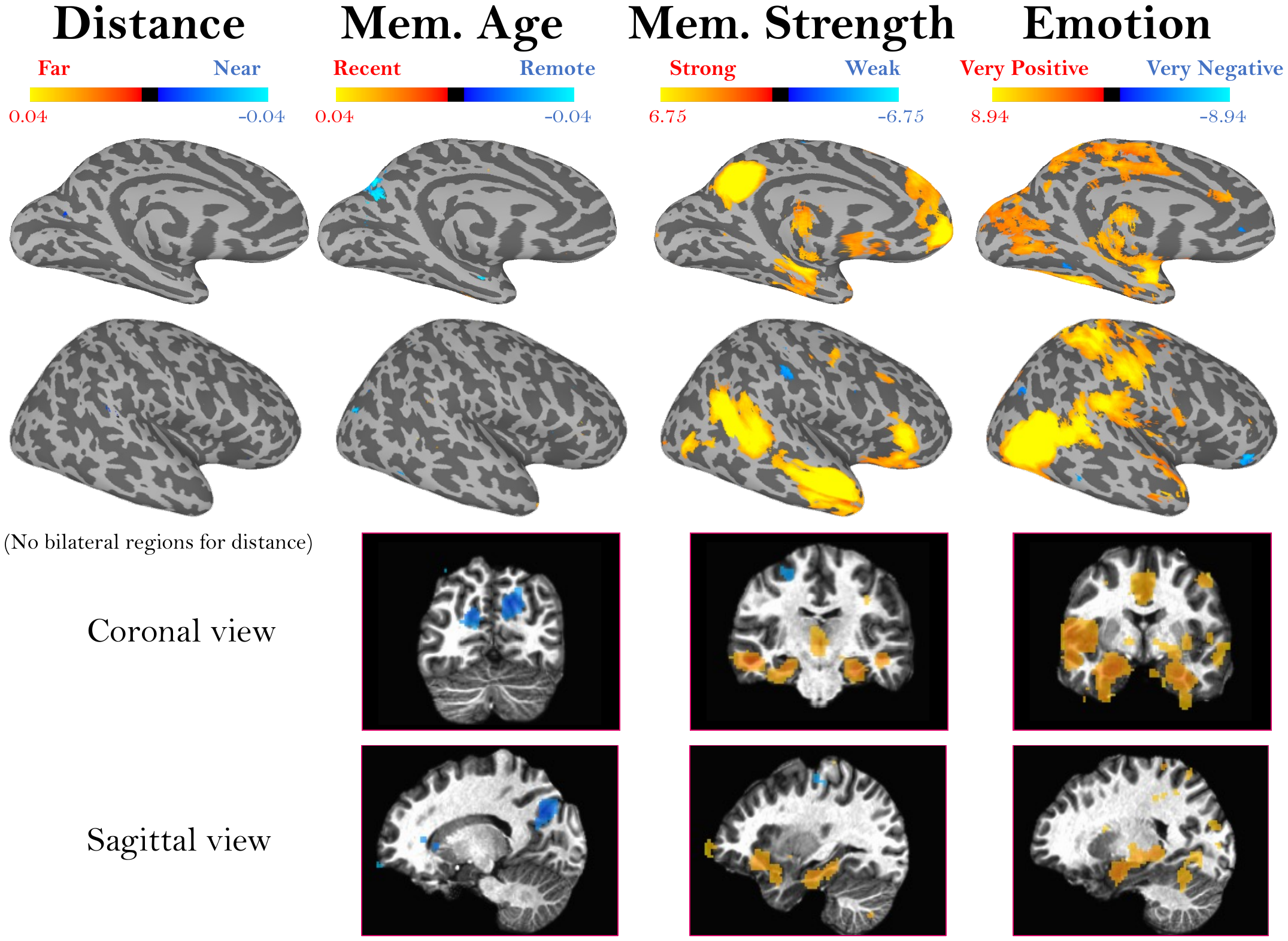

**Supplemental Figure 5.** **Alternate views of memory content representations.** These maps show alternate viewpoints of the results shown in Figure 6. Specifically, these maps show whole-brain results from a multiple regression predicting voxel beta values from separate predictors for a memory’s distance, age, strength, and emotion. Activation represents the mean regressor slope (β) for each predictor, where significance was assessed by comparing the slope across all participants versus a null hypothesis slope of 0 (p < 0.01, uncorrected). The colormaps represent the range of beta values for each predictor, after centering. Shown here are the left medial view, right lateral view, and volumetric coronal and sagittal views. No significant voxels emerged for memory distance (so its blank volumetric views are not shown).

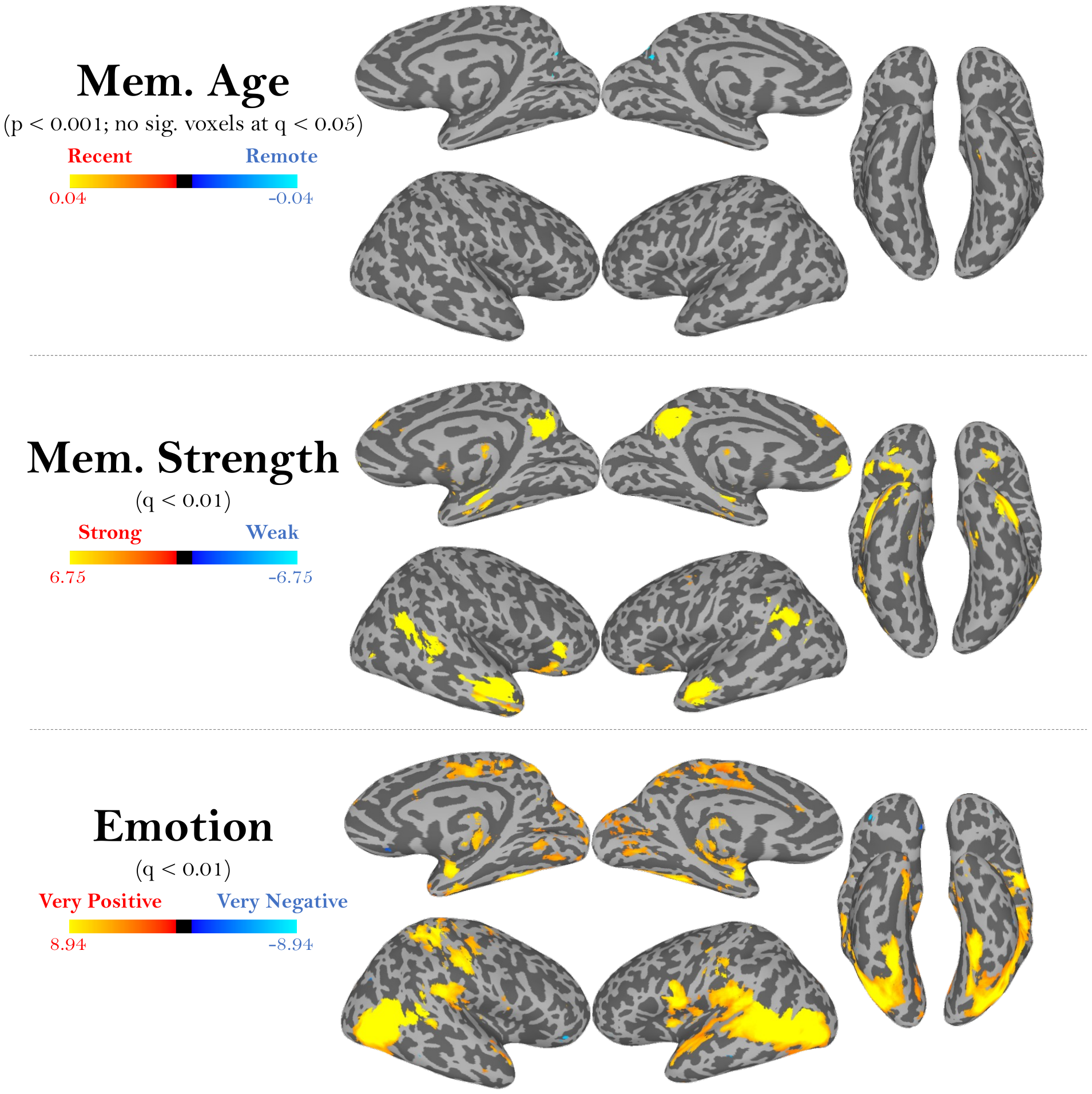
**Supplemental Figure 6.** **Representations of different memory content, at stringent thresholds.** Shown are bilateral maps of the data shown in Figure 6 and Supplemental Figure 5 at more stringent thresholds. These maps show the results from a multiple regression predicting voxel beta values using predictors for a memory’s distance, age, strength and emotion. The colormaps represent the range of beta values for each predictor, after centering. Memory strength and emotion maps are thresholded here at an FDR-corrected threshold of *q* < 0.01. For memory age, no voxels were significant after FDR correction; the above map shows an alternate threshold of *p* < 0.001 uncorrected. No significant voxels emerged for memory distance at either threshold.

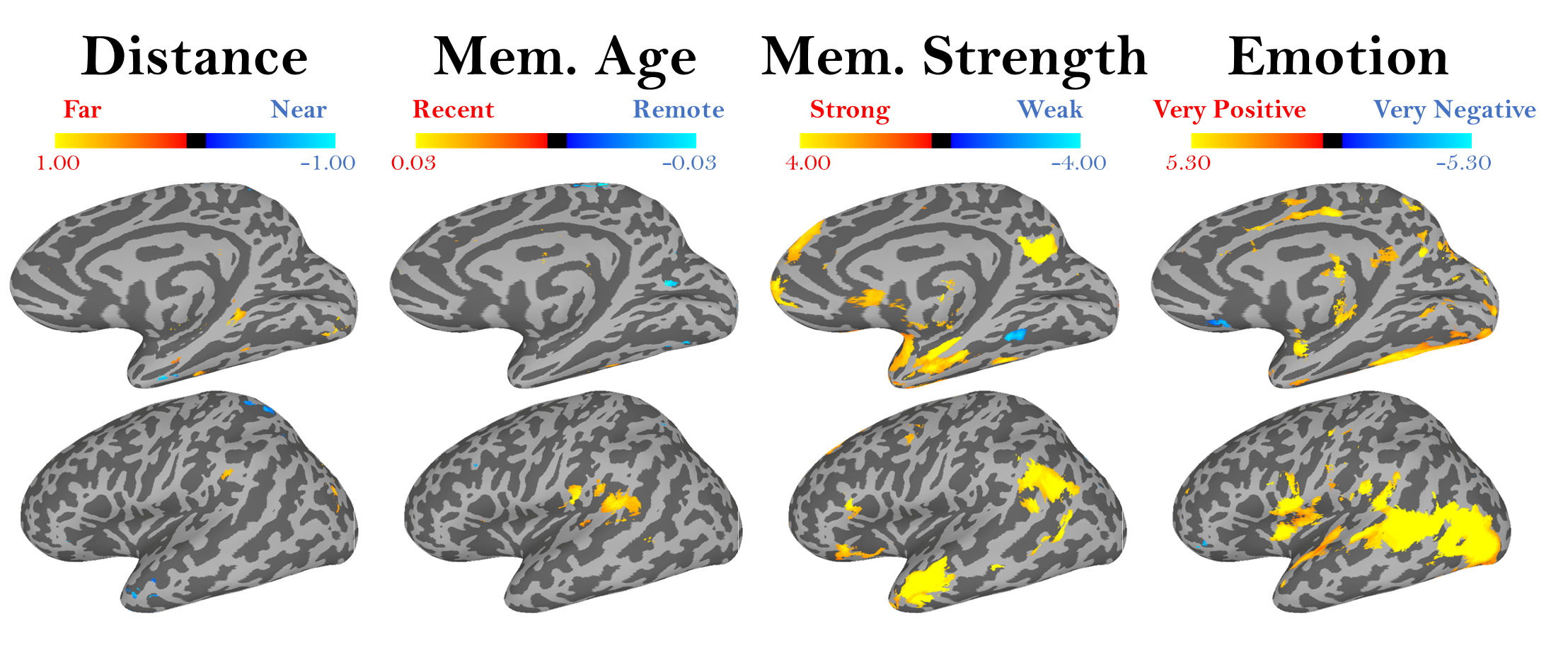
**Supplemental Figure 7.** **Map of memory content representations, for a constrained set of distances (within 50km).** These maps show the same multiple regression as in Figure 6, for predicting voxel beta values from separate predictors for a memory’s distance, age, strength, and emotion. Activation represents the mean regressor slope (β) for each predictor, where significance was assessed by comparing the slope across all participants versus a null hypothesis slope of 0 (p < 0.01, uncorrected). However, in contrast with Figure 6 which used all memory videos, this regression was only conducted with memory videos that occurred within 50km of the scan center, to more closely approximate the short distances used in prior autobiographical memory neuroimaging work (Nielson et al., 2015). 50km was chosen as the optimal distance to cover the broader Washington DC metropolitan area where participants resided, including Northern Virginia and Baltimore. However, this cut-off does eliminate over one-third of the videos recorded in the study (M=111.9 videos, SD=86.4 per participant occurred over 50km away), so statistical power is likely much lower. In these surface maps, the same regions emerge for memory age, memory strength, and emotion as seen when using all memories (Figure 6), such as distinct medial parietal areas sensitive to temporally remote memories and strong memories. New regions emerge sensitive to memory distance along the temporal lobe. However, we find no regions sensitive to distance in the medial parietal cortex, and the temporal regions show no overlap with our anatomically defined medial temporal lobe areas, such as the hippocampus or parahippocampal cortex.

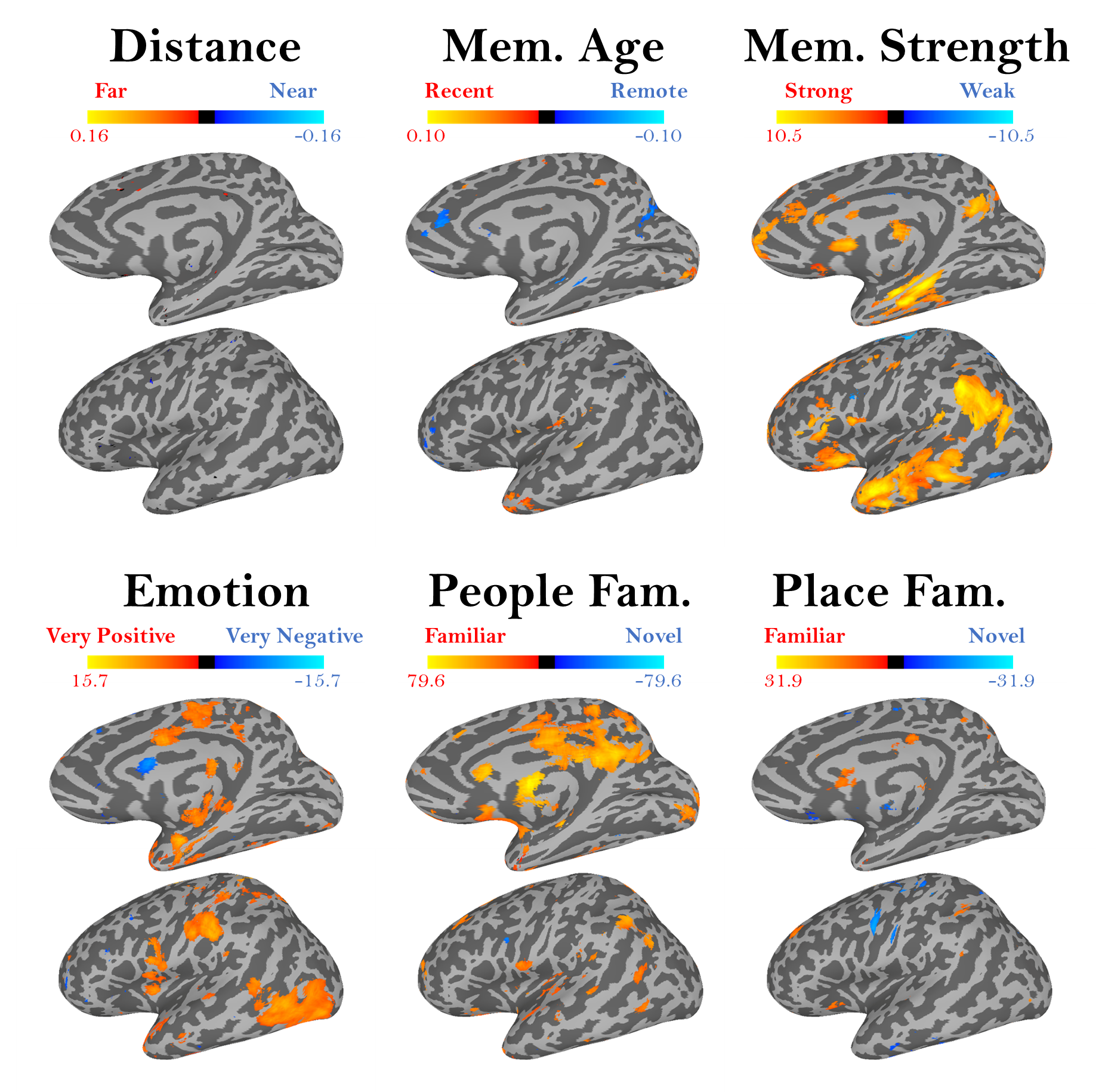
**Supplemental Figure 8.** **Map of memory content representations, with regressors included for people and place familiarity.** These maps show a multiple regression (similar to that in Figure 6), using all memory videos, for predicting voxel beta values from separate predictors for a memory’s distance, age, strength, emotion, people familiarity, and place familiarity. People and place familiarity were modeled categorically. Activation represents the mean regressor slope (β) for each predictor, where significance was assessed by comparing the slope across all participants versus a null hypothesis slope of 0. Given the exploratory nature of this regression, maps are shown at a liberal threshold (p<0.05, uncorrected). Similar regions emerge for memory age, strength, and emotion as in the original regression (Figure 6), including medial parietal regions for memory age and strength. No regions emerge for memory distance. People and place familiarity show sensitivity across medial and lateral parietal regions.

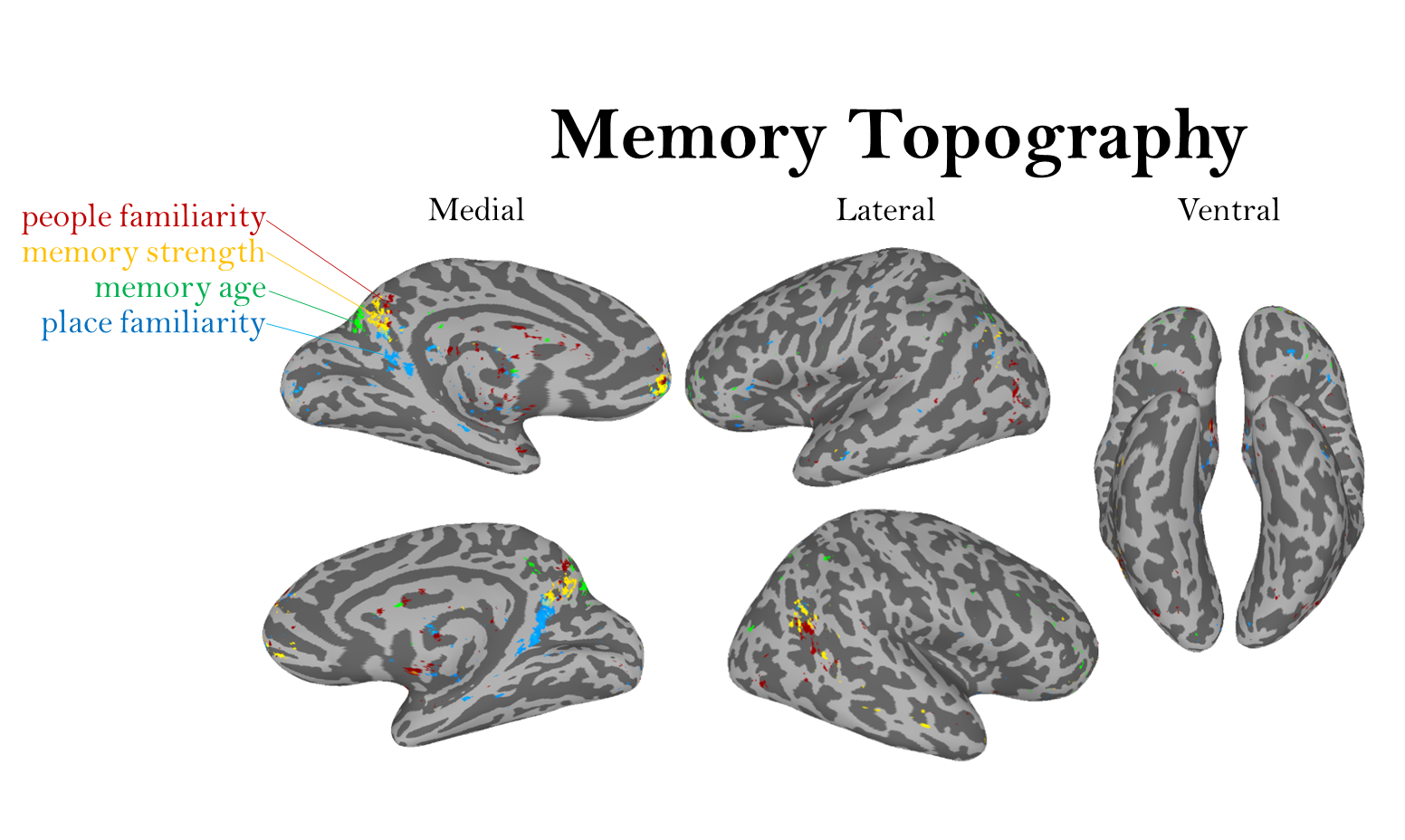

**Supplemental Figure 9.** **Whole-brain views of the memory topography.** These maps show the same information in Figure 8, but across the whole brain. Shown are the top 1000 voxels across the cortex with signal for four different types of information: people familiarity, memory strength, memory age, and place familiarity. Each map is shown at 50% transparency; if a voxel is shared across multiple content types, it will be colored by both maps. While the medial parietal lobe shows the densest distribution of all four types of information with distinct voxel clusters, overlapping regions for people familiarity and memory strength are present in lateral occipital regions. Age, memory strength, and people familiarity also show overlapping sensitivity in orbitofrontal regions.

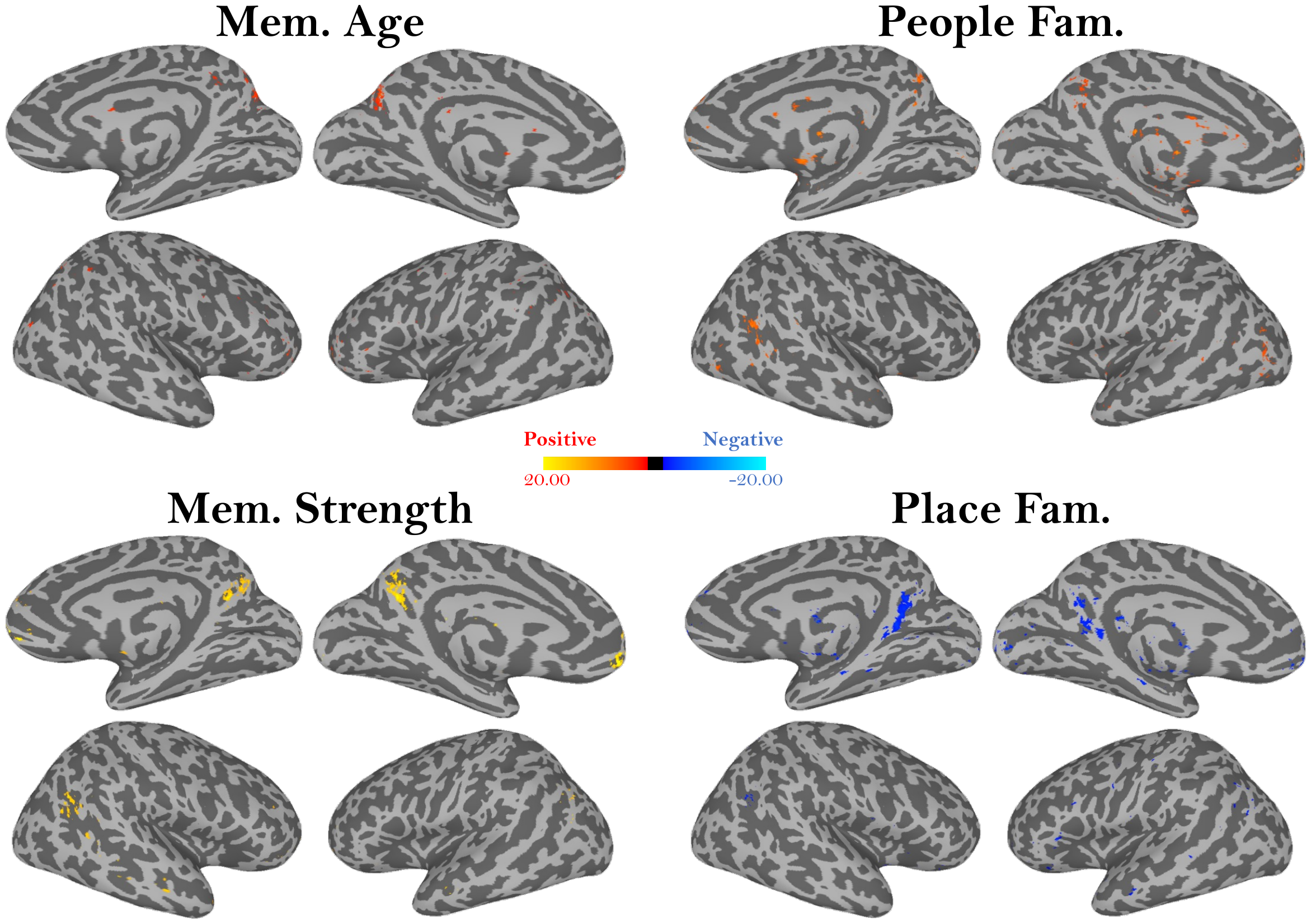

**Supplemental Figure 10.** **Whole-brain views of the memory topography on the same colormap.** These maps show the same data as in Supplemental Figure 9 (the top 1000 voxels for each memory information type), but each information type is on its own brain, and all maps are using the same colormap, representing the range of beta values from -20.00 to 20.00.

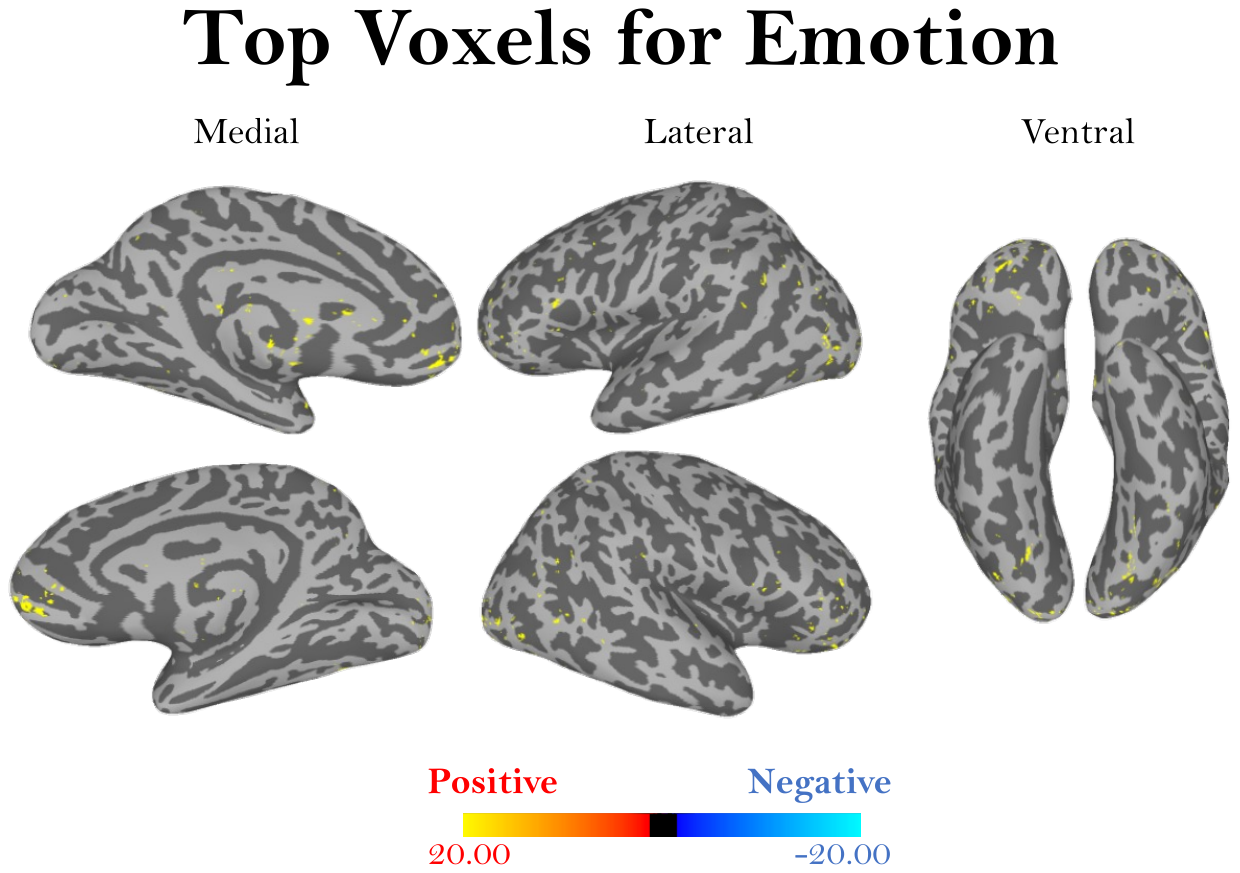

**Supplemental Figure 11.** **Whole-brain views of the top voxels for emotion.** Shown are the top 1000 voxels across the cortex with signal for emotion information in a memory. The colormap shows the range of beta values, on the same range as Supplemental Figure 10. While these peaks for emotion occur in early visual cortex, subcortical areas, and frontal cortex, no voxels appear in the medial parietal cortex.

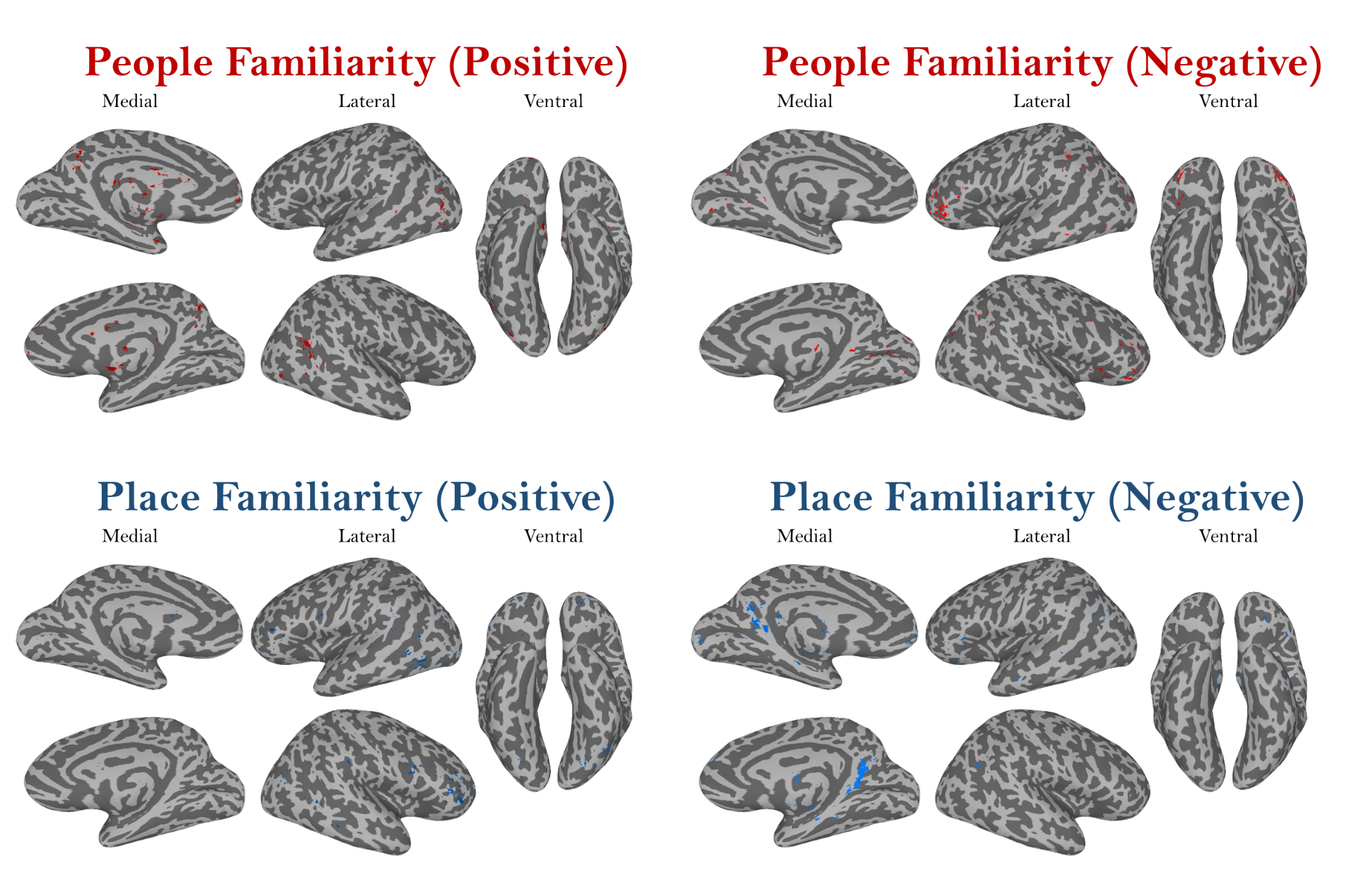

**Supplemental Figure 12.** **Top voxels for people and place familiarity.** These maps show the top 1000 positive and negative voxels (all *p*<0.05) across the cortex for people familiarity (top) and place familiarity (bottom). For people familiarity, orbitofrontal and lateral parietal areas show negative activation, while medial parietal and occipital areas show positive activation. For places, medial parietal areas show the highest negative activation, while orbitofrontal areas show the highest positive activation. Within the topographic distribution of memory content in Figure 8, the medial parietal cortex specifically shows positive activation for people familiarity but negative activation for place familiarity.
